# Structural and mechanistic insights into the Artemis endonuclease and strategies for its inhibition

**DOI:** 10.1101/2021.01.08.423993

**Authors:** Yuliana Yosaatmadja, Hannah T Baddock, Joseph A Newman, Marcin Bielinski, Angeline E Gavard, Shubhashish M M Mukhopadhyay, Adam A Dannerfjord, Christopher J Schofield, Peter J McHugh, Opher Gileadi

## Abstract

Artemis (DCLRE1C) is an endonuclease that plays a key role in development of B- and T-lymphocytes and in DNA double-strand break repair by non-homologous end-joining (NHEJ). Artemis is phosphorylated by DNA-PKcs and acts to open DNA hairpin intermediates generated during V(D)J and class-switch recombination. Consistently, Artemis deficiency leads to radiosensitive congenital severe immune deficiency (RS-SCID). Artemis belongs to a structural superfamily of nucleases that contain conserved metallo-β-lactamase (MBL) and β-CASP (CPSF-Artemis-SNM1-Pso2) domains. Here, we present crystal structures of the catalytic domain of wild type and variant forms of Artemis that cause RS-SCID Omenn syndrome. The truncated catalytic domain of the Artemis is a constitutively active enzyme that with similar activity to a phosphorylated full-length protein. Our structures help explain the basis of the predominantly endonucleolytic activity of Artemis, which contrast with the predominantly exonuclease activity of the closely related SNM1A and SNM1B nucleases. The structures also reveal a second metal binding site in its β-CASP domain that is unique to Artemis. By combining our structural data that from a recently reported structure we were able model the interaction of Artemis with DNA substrates. Moreover, co-crystal structures with inhibitors indicate the potential for structure-guided development of inhibitors.

## INTRODUCTION

Nucleases hydrolyse the phosphodiester bonds of nucleic acids and are grouped into two broad classes: exonucleases and endonucleases. Exonucleases are often non sequence-specific, while endonucleases can be further grouped into sequence-specific endonucleases, such as restriction enzymes, and structure-selective endonucleases [1]. Artemis (also known as SNM1C or DCLRE1C), along with SNM1A (DCLRE1A) and Apollo (SNM1B or DCLRE1B), are human nucleases that fall into the extended structural family of metallo-β-lactamase (MBL) fold enzymes [2,3]. The N-terminal region of Artemis is predicted to have a core MBL fold (aa 1–155, 385–361) with an inserted β-CASP (<u>C</u>PSF73, <u>A</u>rtemis, <u>S</u>NM1 and <u>P</u>SO2) domain (aa 156–384). β-CASP domains are present within the larger family of eukaryotic nucleic acid processing MBLs and confer both DNA/RNA binding and nuclease activity [2]. The C-terminal region of Artemis mediates protein-protein interactions, contains post translational modification (PTM) sites, directs subcellular localisation, and may modulate catalytic activity [4–7].

Although SNM1A, SNM1B, and Artemis are predicted to have similar core structures for their catalytic domains, each have distinct cellular functions and substrate specificities. While SNM1A and SNM1B/Apollo are exclusively 5’ to 3’ exonucleases, the predominant activity of Artemis is endonucleolytic [5,7], although a minor 5’ to 3’ exonuclease activity has been reported [8]. Human SNM1A localises to sites of DNA damage, can digest past DNA damage lesions *in vitro*, and is involved in the repair of interstrand crosslinks (ICLs) [9–11]. SNM1B/Apollo is a shelterin-associated protein required for resection at newly-replicated leading-strand telomeres to generate the 3’-overhang necessary for telomere loop (t-loop) formation and telomere protection [12–14]. Both SNM1A and SNM1B/Apollo prefer ssDNA substrates *in vitro*, with an absolute requirement for a free 5’-phosphate [3,15]. By contrast, Artemis prefers hairpins and DNA junctions as substrates for its endonuclease activity, although it is able to process ssDNA substrates [16–18]

The endonuclease activity of Artemis is responsible for hairpin opening in variable (diversity) joining (V(D)J) recombination [19] and contributes to end-processing in the canonical non-homologous end joining (c-NHEJ) DNA repair [20–23]. V(D)J recombination is initiated by the recognition and binding of recombination-activating gene proteins (RAG1 and RAG2) to the recombination signal sequences (RSSs) adjacent to the V, D, and J gene segments. Upon binding, the RAG proteins induce double-strand breaks (DSBs) and create a hairpin at the coding ends [24–26]. The Ku heterodimer recognises the DNA double-strand break and recruits DNA-dependent protein kinase catalytic subunit (DNA-PKcs) and Artemis to mediate hairpin opening [17]. Following hairpin opening, the NHEJ machinery containing the XRCC4/XLF(PAXX)/DNA-Ligase IV complex is recruited to catalyse the processing and ligation reactions of the DNA ends [20,27,28]. V(D)J recombination is an essential process in antibody maturation [16,29,30]. Mutations in the Artemis gene cause aberrant hairpin opening resulting in severe combined immune deficiency (RS-SCID), with sensitivity to ionising radiation due to impairment of the predominant DSB repair pathway in mammalian cells, NHEJ [17,19,27], and another form of SCID (Omenn syndrome) associated with hypomorphic Artemis mutations [31,32]

One of the most common mutations leading to Artemis loss-of-function are large deletions in the first four exons and a nonsense founder mutation, as found in Navajo and Apache Native Americans [33]. In addition, missense mutations and in-frame deletions in the highly conserved residues such as H35, D165 and H228 can also abolish Artemis’ protein function [34].

Owing to the key roles of Artemis and related DSB repair enzymes in both programmed V(D)J recombination and non-programmed c-NHEJ DSB repair, they are attractive pharmacological targets for the radiosensitisation of tumours.

Here, we present a high-resolution crystal structure of the catalytic core of Artemis (aa1–361) containing both MBL and a β-CASP domains. This reveals that Artemis possesses a unique feature, that is not present in SNM1A and SNM1B/Apollo, i.e., a second metal binding site in its β-CASP domain that bears a resemblance to classical Cys_2_His_2_ zinc finger motifs. We propose that this second metal coordination site is involved in Artemis stabilisation and substrate specificity. We also present a model for Artemis DNA binding based on our data and another recently published structure. The Artemis DNA model is compared with models of DNA binding from related nucleases to reveal distinct features that define a role for Artemis in the end-joining reaction. Following development of an assay suitable for inhibitor screens, we identified drug-like molecules that could potentially inhibit both the Artemis active site and its essential zinc finger-like motif.

## MATERIAL AND METHODS

### Cloning and site directed mutagenesis of WT and mutant Artemis (aa 1-362)

The Artemis MBL-β-CASP domain (WT and mutant) encoding constructs were cloned into the baculovirus expression vector pBF-6HZB which combines an N-terminal His_6_ sequence and the Z-basic tag (GenBank™ accession number KP233213.1) for efficient purification and to promote solubility. The Artemis gene was cloned using ligation independent cloning (LIC) [35]. Site directed mutagenesis was carried out using an inverse PCR experiment whereby an entire plasmid is amplified using complementary mutagenic primers (oligonucleotides) with minimal cloning steps [36]. Using the high-fidelity and high-processivity enzyme Herculase II Fusion DNA Polymerase (Agilent), a PCR was performed to amplify a whole plasmid. The PCR product was then added to a KLD enzyme mix (NEB) reaction and was incubated at room temperature for 1 hour, prior to transformation into *Escherichia coli* cells.

### Expression and purification of WT and mutant Artemis with IMAC (aa 1-362)

Baculovirus generation was performed as previously described [3]. Recombinant proteins were produced in *Sf9* cell at 2 × 10^6^ cells/ mL infected with 1.5 mL of P2 virus for WT and 3 mL of P2 virus for mutants respectively. Infected *Sf9* cells were harvested 70 h after infection by centrifugation (900 × g, 20 min). The cell pellet was resuspended in 30 mL/ L lysis buffer (50 mM HEPES pH 7.5, 500 mM NaCl, 10 mM imidazole, 5% (v/v) glycerol and 1 mM TCEP), snap frozen in liquid nitrogen, then stored at −80 °C for later use.

Thawed cell aliquots were lysed by sonication. The lysates were clarified by centrifugation (40,000 g, 30 min), then the supernatant was passed through a 0.80 μm filter (Millipore) and loaded onto an equilibrated (lysis buffer) immobilised metal affinity chromatography column (IMAC) (Ni-NTA Superflow Cartridge, Qiagen). The immobilised protein was washed with lysis buffer, then eluted using a linear gradient of elution buffer (50 mM HEPES pH 7.5, 500 mM NaCl, 300 mM imidazole, 5% v/v glycerol, and 1 mM TCEP). The protein containing fractions were pooled and passed through an ion exchange column (HiTrap^®^ SP FF GE Healthcare Life Sciences) pre-equilibrated in the SP buffer A (25 mM HEPES pH 7.5, 300 mM NaCl, 5% (v/v) glycerol and 1 mM TCEP). The protein was eluted using a linear gradient of SP buffer B (SP buffer A with 1 M NaCl), and fractions containing the tag-free Artemis were identified by electrophoresis.

Artemis containing fractions were pooled and dialysed overnight at 4°C in SP buffer A and supplemented with recombinant tobacco etch virus (TEV) protease for cleavage of the _6_His-ZB tag. The protein was subsequently loaded into an ion exchange column (HiTrap^®^ SP FF GE Healthcare Life Sciences), pre-equilibrated in the SP buffer A to remove _6_His-ZB tag and uncleaved protein. The protein was eluted using a linear gradient of SP buffer B, and fractions containing the tag-free Artemis were identified by electrophoresis. Artemis-containing fractions from the SP column elution were combined and concentrated to 1 mL using a 30 kDa MWCO centrifugal concentrator. The protein was then loaded on to a Superdex 75 increase 10/300 GL equilibrated with SEC buffer (25 mM HEPES pH 7.5, 300 mM NaCl, 5% (v/v) glycerol, 2 mM TCEP).

Mass spectrometric analysis of the purified proteins revealed masses of 41716.5 Da, 41650.5 Da, 41672.2 Da, 41639.9 Da for WT, H35A, D37A and H35D proteins, respectively. The calculated masses are 41715.09, 41649.2, 41671.2 and 14639.2, respectively, all within 1.5 Da of the measured masses.

### Expression and purification of WT truncated Artemis catalytic domain without IMAC (aa 1-362)

The truncated Artemis protein was expressed and purified in a similar manner as described above except for the first purification step. We used 5 mL HiTrap^®^ SP Fast Flow (GE Health Care) column as the first step of purification. Following an overnight TEV cleavage the protein was subjected to a second ion exchange step (5 mL HiTrap^®^ SP Fast Flow (GE Health Care)) for the removal of the Z-Basic protein tag. The protein was further purified by size exclusion chromatography (Highload^®^ 16/200 Superdex^®^ 200).

### Cloning, expression and purification of full-length WT Artemis (aa 1-692)

The full-length Artemis encoding construct was cloned into pFB-CT10HF-LIC, a baculovirus expression vector containing a C-terminal His_10_ and FLAG tag. pFB-CT10HF-LIC was a gift from Nicola Burgess-Brown (Addgene plasmid # 39191; http://n2t.net/addgene:39191; RRID: Addgene_39191). As for the truncated protein, the full-length Artemis gene was also cloned using ligation independent cloning (LIC) [35].

The baculovirus mediated expression of the full length DCLRE1C/ Artemis gene was performed in a manner similar to that used for the truncated protein. However, instead of infection with 1.5 mL of P2 Virus, 3.0 mL of P2 virus was used to infect *Sf9* cells at 2 × 10^6^ cells/ mL for the expression of the full-length Artemis construct.

Cell harvesting and the initial IMAC purification steps were performed as described for the catalytic domain. Following IMAC chromatographic purification, TEV cleavage overnight in dialysis buffer (50 mM HEPES pH 7.5, 0.5 M NaCl, 5% glycerol and 1 mM TCEP) gave protein which was then passed through a 5 mL Ni-sepharose column; the flowthrough fractions were collected. The Artemis protein was then concentrated using a centrifugal concentrator (Centricon, MWCO 30 kDa) before loading on a Superdex S200 HR 16/60 gel filtration column in dialysis buffer. Fractions containing purified Artemis protein were pooled and concentrated to 10 mg/mL.

### Electospray mass spectrometry (ESI-QTOF)

Reversed-phase chromatography was performed in-line prior to mass spectrometry using an Agilent 1290 uHPLC system (Agilent Technologies inc. – Palo Alto, CA, USA). Concentrated protein samples were diluted to 0.02 mg/ml in 0.1% formic acid and 50 μl was injected on to a 2.1 mm × 12.5 mm *Zorbax* 5um 300SB-C3 guard column housed in a column oven set at 40 °C. The solvent system used consisted of 0.1% formic acid in ultra-high purity water (Millipore) (solvent A) and 0.1 % formic acid in methanol (LC-MS grade, Chromasolve) (solvent B). Chromatography was performed as follows: Initial conditions were 90 % A and 10 % B and a flow rate of 1.0 ml/min. A linear gradient from 10 % B to 80 % B was applied over 35 seconds. Elution then proceeded isocratically at 95 % B for 40 seconds followed by equilibration at initial conditions for a further 15 seconds. Protein intact mass was determined using a 6530 electrospray ionisation quadrupole time-of-flight mass spectrometer (Agilent Technologies Inc. – Palo Alto, CA, USA). The instrument was configured with the standard ESI source and operated in positive ion mode. The ion source was operated with the capillary voltage at 4000 V, nebulizer pressure at 60 psig, drying gas at 350°C and drying gas flow rate at 12 L/min. The instrument ion optic voltages were as follows: fragmentor 250 V, skimmer 60 V and octopole RF 250 V.

### Protein crystallisation and Soaking

Artemis (PDB:6TT5) was crystallised using the sitting drop vapour diffusion method by mixing 50 nL protein with 50 nL crystallisation solution comprising 0.2 M ammonium chloride, 20% (v/v) PEG 3350. Crystals grew after 2 weeks and reached maximum size within 3 weeks. An unliganded crystal was flash frozen in liquid nitrogen, cryoprotected with the mother liquor supplemented with 20% (v/v) ethylene glycol solution.

The non-IMAC purified Artemis (PDB:7AF1) was crystallised in a similar manner, with the addition of 20 nL of crystal seed solution obtained from previous crystallisation experiment. The crystals were grown in a solution comprising 0.25 M ammonium chloride and 30% (v/v) PEG 3350 at 4°C. Crystals grew after one day and reached a maximum size within one week.

Artemis variants (mutants H33A and H35D) were crystallised using the sitting drop vapour diffusion method by mixing 50 nL protein with 50 nL crystallisation solution comprising 0.1 M sodium citrate pH 5.5, 20% PEG 3350, while the D37A was crystalised in 0.2 M ammonium acetate, 0.1 M bis-TRIS pH 5.5, 25% PEG 3350. All Artemis variants were crystalised in the presence of 20 nL of crystal seed solution obtained from previous crystallisation experiment. Crystals grew after one day at 4°C. and reached maximum size within one week

### Data collection and refinements

Data were collected at Diamond Light Source I04, I03, or I24 beamlines. Diffraction data were processed using DIALS [37] and structures were solved by molecular replacement using PHASER [38] and the PDB coordinates 5Q2A. Model building and the addition of water molecules were performed in COOT [39] and structures refined using REFMAC [40]. Data collection and refinement statistics are given in Table I. The X-ray fluorescence data was collected at Diamond Light Source I03 (6TT5) using 100% transmission and 12.7 eV, and I24 (7AF1) using 1% transmission and 12.8 eV (Suppl. Figure 1).

### Generation of 3’-radiolabelled substrates

10 pmol of single-stranded DNA (Eurofins MWG Operon, Germany) were labelled with 3.3 pmol of α-^32^P-dATP (Perkin Elmer) by incubation with terminal deoxynucleotidyl transferase (TdT, 20 U; ThermoFisher Scientific), at 37°C for 1 hour. This solution was then passed through a P6 Micro Bio-Spin chromatography column (BioRad), and the radiolabeled DNA was annealed with the appropriate unlabeled oligonucleotides (1:1.5 molar ratio of labelled to unlabeled oligonucleotide) (Supplementary Table 1 for sequences) by heating to 95°C for 5 min, and cooling to below 30°C in annealing buffer (10 mM Tris-HCl; pH 7.5, 100 mM NaCl, 0.1 mM EDTA).

### Gel-based nuclease assays

Standard nuclease assays were carried out in reactions containing 20 mM HEPES-KOH, pH 7.5, 50 mM KCl, 10 mM MgCl_2_, 0.05% (v/v) Triton X-100, 5% (v/v) glycerol (final volume: 10 μL), and the indicated concentrations of Artemis. Reactions were started by the addition of DNA substrate (10 nM), incubated at 37°C for the indicated time, then quenched by addition of 10 μL stop solution (95% formamide, 10 mM EDTA, 0.25% (v/v) xylene cyanole, 0.25% (v/v) bromophenol blue) with incubation at 95 °C for 3 min.

Reaction products were analysed by 20% denaturing polyacrylamide gel electrophoresis (made from 40% solution of 19:1 acrylamide:bis-acrylamide, BioRad) and 7 M urea (Sigma Aldrich)) in 1 x TBE (Tris-borate EDTA) buffer. Electrophoresis was carried out at 700 V for 75 minutes; gels were subsequently fixed for 40 minutes in a 50% methanol, 10% acetic acid solution, and dried at 80°C for two hours under a vacuum. Dried gels were exposed to a Kodak phosphor imager screen and scanned using a Typhoon 9500 instrument (GE).

### Fluorescence-based nuclease assay

The protocol of Lee *et al* [41] was adapted for structure-specific endonuclease activity. A ssDNA substrate was utilised containing a 5’ FITC-conjugated T and a 3’ BHQ-1 (black hole quencher)-conjugated T (Suppl. Table 1). As the FITC and BHQ-1 are located proximal to one another, prior to endonucleolytic incision, the intact substrate does not fluoresce. Following endonucleolytic incision by DCLRE1C/Artemis, there is uncoupling of the FITC from the BHQ-1 and an increase in fluorescence. Inhibitors (at increasing concentrations) were incubated with Artemis for 10 minutes at room temperature, before the reaction was started with the addition of DNA substrate. Assays were carried out in a 384-well format, in a 25 μL reaction volume. The buffer was the same as for the gel-based nuclease assays, Artemis concentration was 50 nM, and the DNA substrate was at 25 nM. Fluorescence spectra were measured using a PHERAstar FSX (excitation: 495 nm; emission: 525 nm) with readings taken every 150 sec, for 35 min, at 37 °C.

## RESULTS

### Human Artemis (SNM1C or DCLRE1C) has a core catalytic fold similar to SNM1A and Apollo/SNM1B

The core catalytic domain of Artemis (aa 3–361) was produced in baculovirus-infected *Sf9* cells fused to a highly basic His_6_-Zb tag, which confers tight binding to cation exchange columns. The protein was purified using immobilised metal affinity chromatography (IMAC) on a Nickel-Sepharose column as the initial step. Subsequent preparations were performed without the use of IMAC, to avoid the introduction of Ni^2+^ ions during purification.

Artemis protein was purified as detailed in the Methods & Materials, and crystals were subsequently grown and diffracted to 1.6 Å resolution (Table 1); the structure was solved using a structure of SNM1A (PDB coordinates 5Q2A) as the molecular replacement model. The resultant Artemis structure (PDB coordinates 7AF1) contains a single molecule in the asymmetric unit, with two zinc ions coordinated at the active site. The metal ions were identified using X-ray fluorescence (XRF) analysis during data collection at the Diamond Light source. When using the protein purified using IMAC, the first zinc ion in the active site can be replaced by a nickel ion (PDB: 6TT5) (Figure 2E). The presence of the nickel ion was also confirmed in the crystal by XRF. The X-ray fluorescence analysis of the metal ions present in the structures are shown in Suppl. Figure 1. This metal ion coordination pattern has been observed with other member of the family, such as SNM1A and SNM1B/Apollo [3,9].

**Table 1:** Data collections and refinement statistics. * Data in parentheses is for the high-resolution shell.

| PDB ID | 6TT5<br>(Ni and Zn) | 7AF1 (2 Zn) | 7AFS<br>(D37A) | 7AFU<br>(H33A) | 7AGI<br>(H35D) | 7APV<br>(Ceftriaxone) | 7ABS<br>(DNA<br>bound) |
| --- | --- | --- | --- | --- | --- | --- | --- |
| **Data Collection**<br>**and processing** |  |  |  |  |  |  |  |
| Diffraction Source | DLS (I04) | DLS (I24) | DLS (I03) | DLS (I03) | DLS (I03) | DLS (I03) | APS 17-IDD |
| Wavelength (Å) | 0.979 | 0.976 | 0.976 | 0.976 | 0.976 | 0.976 | 1.00 |
| Space group | P1 | P1 | P1 | P1 | P1 | P1 | P21212 |
| Cell Dimensions |  |  |  |  |  |  |  |
| $a, b, c$ (Å) | 35.87,<br>47.99, 48.25 | 35.75,<br>47.97, 48.15 | 35.91,<br>48.06,<br>48.21 | 35.97,<br>47.90,<br>48.37 | 35.97,<br>48.05, 48.44 | 35.88, 48.10,<br>48.25 | 72.81,<br>111.00,<br>55.17 |
| $\alpha, \beta, \gamma$ (°) | 82.61,<br>76.37, 85.98 | 82.68, 76.35,<br>85.81 | 82.89,<br>76.43,<br>86.38 | 82.51,<br>75.94,<br>87.73 | 82.43, 76.01,<br>87.33 | 82.76, 76.29,<br>86.30 | 90.00,<br>90.00,<br>90.00 |
| Resolution (Å) * | 47.55 - 1.50<br>(1.53 - 1.50) | 47.53 - 1.70<br>(1.73 - 1.70) | 35.35 -<br>1.70<br>(1.73 -<br>1.70) | 35.53 -<br>1.56<br>(1.59 -<br>1.56) | 47.62 - 1.70<br>(1.73 - 1.70) | 47.69 - 1.95<br>(2.00 - 1.95) | 40 -<br>1.97<br>(2.02 -<br>1.97) |
| R<sub>merge</sub> (%)* | 6.0 (64.4) | 13.3 (79.5) | 5.3<br>(53.2) | 4.9<br>(31.5) | 4.8 (23.2) | 11.6 (65.0) | 5.0<br>(82.8) |
| $I/\sigma(I)$ | 13.4 (3.2) | 5.8 (2.0) | 11.7<br>(2.2) | 11.3<br>(2.6) | 13.4 (3.8) | 8.6 (2.8) | 16 (2.4) |
| Completeness (%) | 94.7 (66.4) | 97.4 (95.6) | 97.3<br>(96.0) | 94.2<br>(63.7) | 97.4 (95.4) | 98.0 (97.2) | 99.6<br>(99.6) |
| Multiplicity | 3.6 (3.2) | 3.5 (3.5) | 3.6 (3.7) | 3.6 (3.3) | 3.7 (3.8) | 3.5 (3.6) | 6.4 (6.6) |
| **Refinements** |  |  |  |  |  |  |  |
| Resolution (Å) | 47.55 - 1.50 | 47.53 - 1.70 | 47.66 -<br>1.70 | 35.53 -<br>1.56 | 47.62 - 1.70 | 47.96 - 1.95 | 40 - 1.97 |
| No. of reflections | 44746 | 31282 | 34470 | 39478 | 31745 | 21086 | 32168 |
| R<sub>work</sub> | 0.17 | 0.19 | 0.18 | 0.19 | 0.19 | 0.18 | 0.22 |
| R<sub>free</sub> | 0.19 | 0.21 | 0.22 | 0.21 | 0.21 | 0.23 | 0.28 |
| No. of Atom |  |  |  |  |  |  |  |
| Protein | 2927 | 2989 | 2957 | 2985 | 3122 | 2975 | 2923 |
| Water | 224 | 144 | 163 | 255 | 164 | 107 | 105 |
| Zinc/ Nickle | 3 | 3 | 2 | 1 | 1 | 2 | 3 |
| Ethylene glycol | 24 | 28 | 12 | 44 | 32 | 16 | - |
| DNA | - | - | - | - | - | - | 253 |
| Ceftriaxone | - | - | - | - | - | 1 | - |
| B-factors | 16.3 | 19.3 | 25.9 | 17.5 | 18.4 | 22.6 | 56 |
| r.m.s. deviations |  |  |  |  |  |  |  |
| Bond length (Å) | 0.003 | 0.008 | 0.008 | 0.006 | 0.007 | 0.01 | 0.007 |
| bond angles (°) | 1.23 | 1.33 | 1.29 | 1.28 | 1.32 | 1.5 | 1.47 |

The overall fold of human Artemis protein catalytic core is very similar to that of human SNM1A and SNM1B/Apollo (2 Å RMSD). It has the key structural characteristics of human MBL fold nucleases, with the di-metal containing active site interfaced between the MBL and β-CASP domains (Figure 1A). As anticipated, the MBL domain (Figure 1A and 1B, in pink) of Artemis has the typical α/β-β/α sandwich MBL fold [42] and contains all of the highly conserved motifs 1–4 (Figure 1C and D; and sequence alignment, Suppl. Figure 2) which are typical for the whole MBL superfamily, and motif A–C which are typical of the β-CASP fold containing family [2,27,43,44].

**Figure 1:**
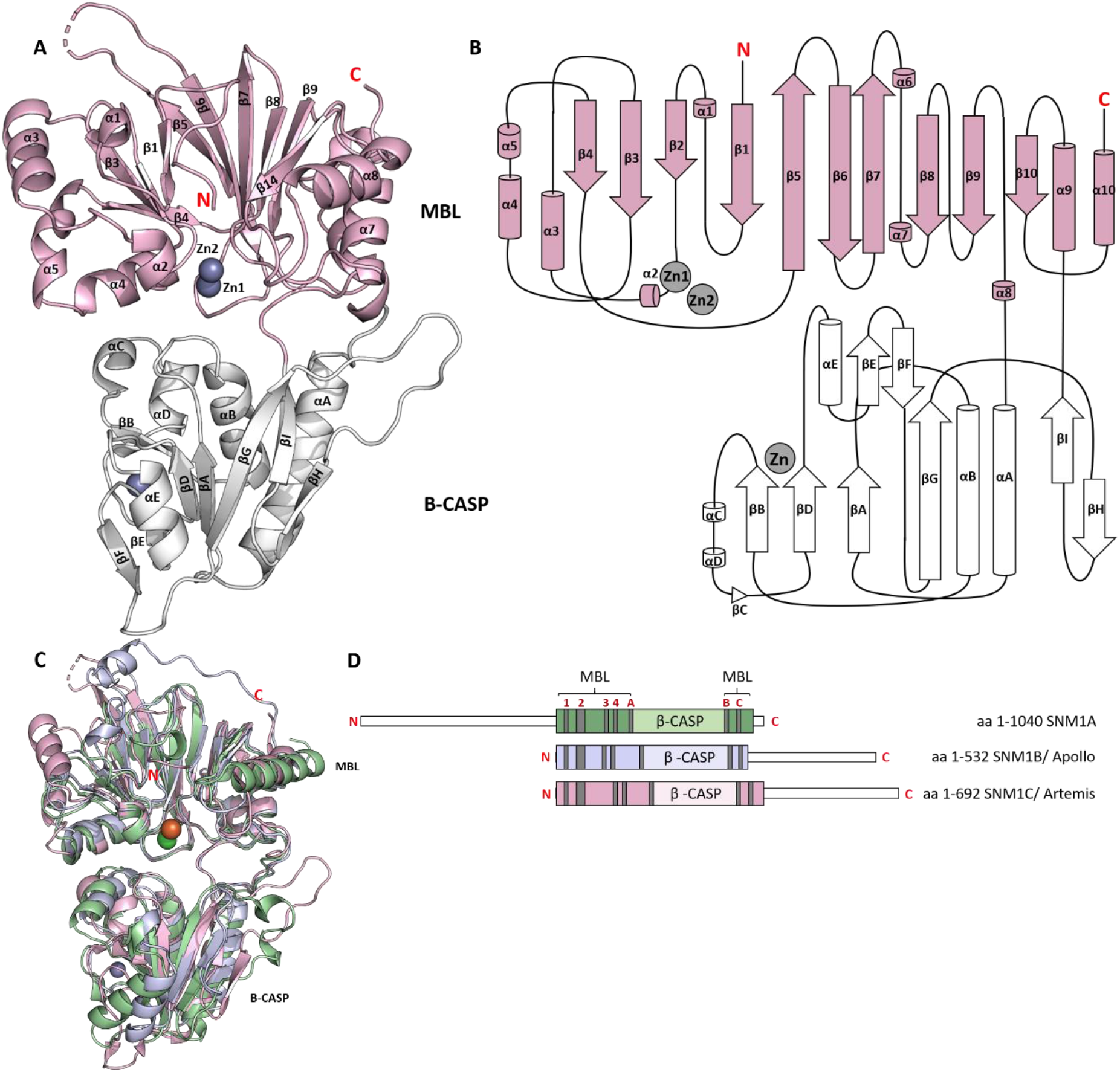
Overall architecture of Artemis. **A.** A cartoon representation of the structure of human SNM1C/ Artemis. The active site containing MBL domain is in pink; the β-CASP domain (white) contains a novel zinc-finger like motif, that has not been identified in other MBL/ β-CASP nucleic acid processing enzymes. The three zinc ions are represented by grey spheres. **B**. Topology diagram of Artemis protein. The β-strands are represented arrows and α-helices by cylinders. The MBL domain (pink) has the typical α/β-β/α sandwich of the MBL superfamily, with an insert of the β-CASP domain (white) between the small helix α6 and helix α7. **C**. Overlay of structures of the human SNM1 Family members: SNM1A, SNM1B and SNM1C. **D.** Cartoon representation of amino acid sequence alignment for human SNM1A, SNM1B and SNM1C, showing the conserved MBL and b-CASP domains.

Motifs 1–4 (Figures 1D and 2C) are responsible for metal ion coordination in both DNA and RNA processing MBL enzymes [44]. As previously observed in crystal structures of human SNM1A and Apollo/SNM1B, Artemis can coordinate one or two metal ions in its active site. One zinc ion (Zn1) in the active site is coordinated by four residues (His33, His35, His115, and Asp116) and two water molecules (H_2_O 506 and 611) in an octahedral manner (Figure 2C). The second zinc ion (Zn2) was refined with 30% occupancy and is coordinated by three residues (Asp37, His38, and Asp136) and two water molecules (H_2_O 506 and 529). The low occupancy of the second zinc ion, together with the two conformations (0.5 occupancy for each conformation) observed for Asp37 (Figure 2E) suggest that this site binds a metal ion less tightly than the Zn1 site, consistent with studies on other human MBL fold nucleases [3, Baddock *et. al.,2020*]

**Figure 2:**
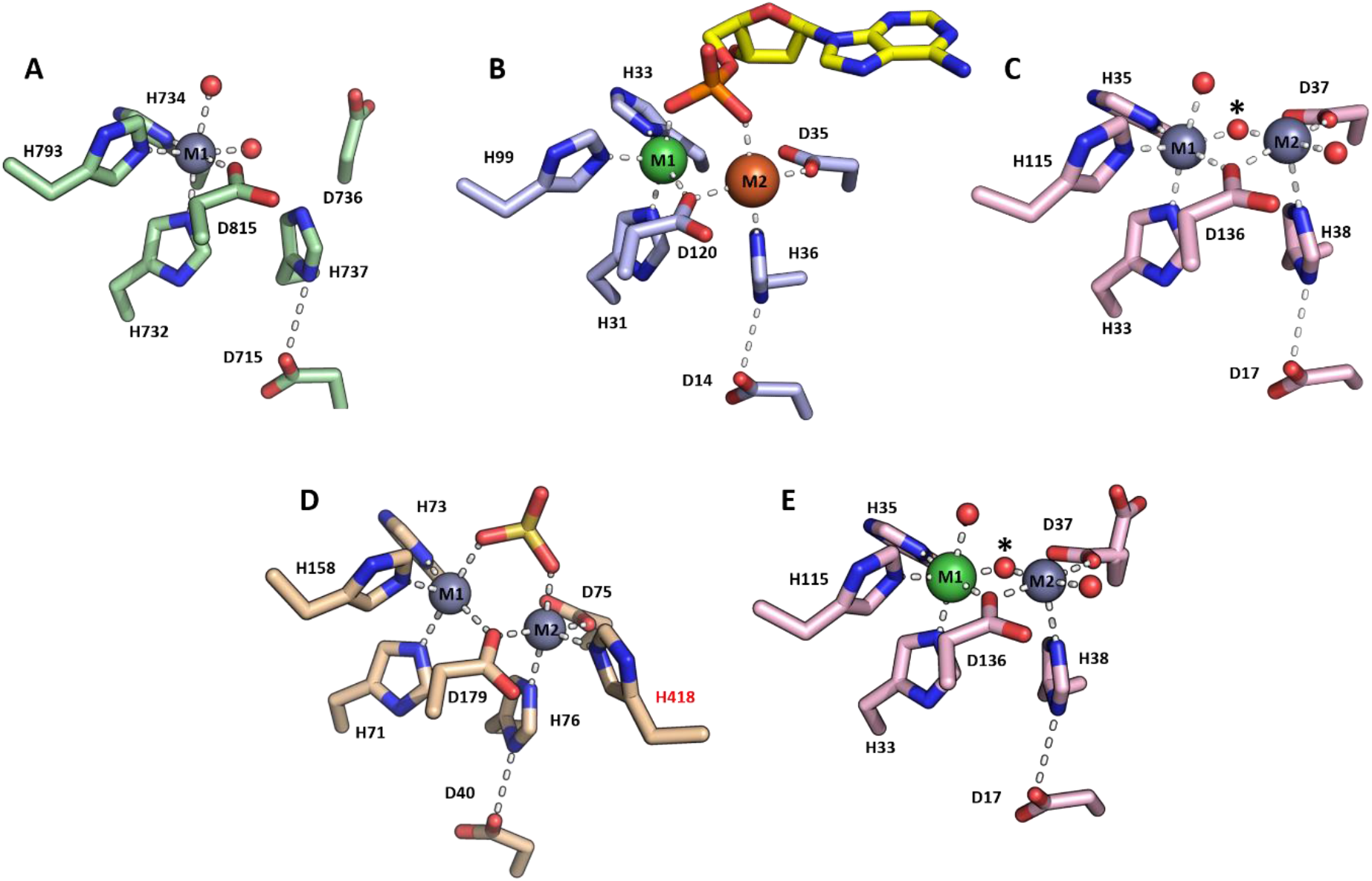
Active site views of human MBL/β-CASP nuclease family enzymes. Each of these catalytic sites contains 4 highly conserved motifs (1-4, in red). Motif 1 = Asp, motif 2 = 3 His and 1 Asp (HxHxDH), motif 3 = His and motif 4 = Asp. **A.** The human SNM1A active site with a single octahedral zinc ion (grey) coordination (PDB: 5AHR). **B**. The active site of human SNM1B/ Apollo (PDB: 7A1F) with a nickel ion (green) and an iron ion (orange) with a coordinating AMP molecule. **C**. Human SNM1C/ Artemis (PDB: 7AF1) purified in the absence of IMAC with two zinc ions (in grey) in its active site. A water molecule shared (asterisk*) between the two metals is the proposed nucleophile for the hydrolytic reaction. **D.** The active site of the human RNA processing enzyme CPSF73 (PDB: 2I7T). The second zinc ion (M2) is coordinated by an additional histidine residue (His 418) which has no counterpart in the SNM1 proteins. **E.** The active site of human SNM1C/ Artemis (PDB: 6TT5) purified with IMAC. A nickel ion is present in the first metal coordination site.

The structure of human SNM1A (PDB: 5AHR) [3] was solved with a single zinc ion coordinated in the active site (Figure 2A). By contrast, SNM1B structures solved with a bound AMP (Baddock *et. al.,* accompanying paper) showed that both metal ions are positioned to coordinate the phosphate group of the AMP in an octahedral manner (Figure 2B). In summary, the octahedral coordination sites for the first zinc ion are contributed to by three histidines, one aspartate residue, and either water molecules or a phosphate oxygen of the substrate; the second metal ion is more weakly coordinated in the SNM1 protein family, with one histidine and two aspartates, with the remaining three positions occupied by water or a phosphate oxygen of the DNA substrate. This can explain the partial occupancy of the second zinc position in SNM1 enzyme structures, where the full occupancy may be achieved only in presence of substrate. We propose that Artemis would coordinate a phosphate group of its substrate in a similar manner. The structure of human CPSF-73 (PDB: 2I7T), an RNA processing nuclease, with two active site bound zinc ions and a phosphate molecule [45], shows that the two zinc ions are coordinated in a very similar geometry with the human MBL DNA processing enzymes. [46,47]. A striking difference between the MBL RNA and DNA nucleases is that the second metal ion (M2) in the RNA processing nucleases is coordinated by an additional histidine residue (His418 for CPSF-73) [45] that is absent in the DNA processing enzymes (Figure 2D).

### The structure of Artemis reveals a novel zinc-finger like motif in the β-CASP domain

Proteins with a β-CASP fold form a distinct sub-group within the MBL-fold superfamily that specifically act on nucleic acids [2]. Artemis’ β-CASP domain is comprised of residues 156–384 and it is the second globular domain (Figure 1A, shown in white) in the catalytic region; inserted within the Artemis MBL fold sequence between the small α-helices 6 and 7 (figure 1 B). The β-CASP domain has been proposed to facilitate substrate recognition and binding in the nucleic acid processing MBL fold-containing family of enzymes [2,48].

Another metal ion coordination site, unique to Artemis, is present in the β-CASP domain, with similarity to the canonical Cys_2_ His_2_ zinc-finger motif [49,50]. Many DNA binding proteins, including transcription factors and a substantial number of DNA repair factors (including those involved in NHEJ), possess the classical Cys_2_ His_2_ zinc finger motif, that serves as a structural feature stabilising the DNA binding domain [49–52]. A typical Cys_2_ His_2_ zinc coordinating finger (Figure 3A) has a ββα motif, wherein the zinc ion is coordinated between an α-helix and two antiparallel β-sheets. The zinc ion confers structural stability and hydrophobic residues located at the sides of the zinc coordination site enable specific binding of the zinc finger in the major groove of the DNA [49,50,53,54]. Similar to the canonical Cys_2_ His_2_ zinc finger motif, the zinc ion coordination in Artemis’ β-CASP domain adopts a tetrahedral geometry, with coordination by two cysteine (Cys256 and Cys272) and two histidine (His228 and His254) residues (Figure 3 B). However, in the case of Artemis the metal ion coordination site is sandwiched between two β-sheets instead of an α-helix and two antiparallel β-sheets.

**Figure 3:**
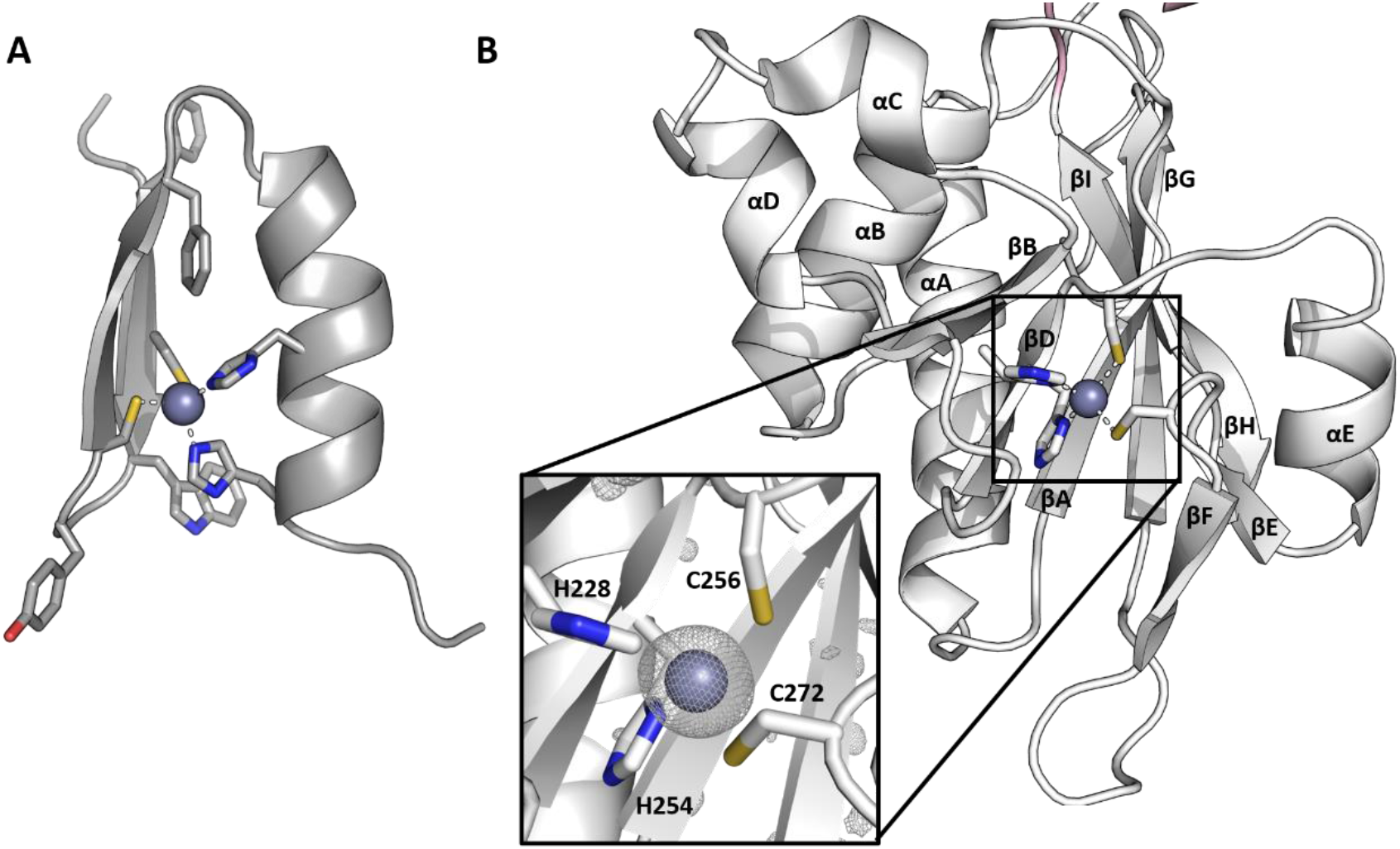
Comparison of a novel zinc-finger like motif in the β-CASP domain of Artemis with a canonical zinc-finger motif. **A.** Cartoon representation of the classical Cys_2_His_2_ zinc-finger motif, from transcription factor SP1F2 (PDB code: 1SP2). This has a ββα fold, where two Cys- and two His-residues are involved in zinc ion coordination and the sidechains of three conserved hydrophobic residues are shown. **B.** The β-CASP region of Artemis has a novel zinc-finger like motif. The inset shows the four residues (two His and two Cys) coordinating the zinc ion (grey). The *F_o_* – *F_c_* electron-density map (scaled to 2.5σ in PyMOL) surrounding the zinc ion before it was included in refinement.

Almost all the residues in the zinc-finger like motif (His228, Cys256, and Cys227) are unique to Artemis (sequence alignment Suppl. Figure 2), with only His254 being well conserved within the SNM1-family. However, these four residues that forms the zinc-finger like motif are highly conserved in Artemis across different species (from human to marine sponge), implying functional importance (sequence alignment Suppl. Figure 3). Consistently, substitution of His228 and His254 (H228N and H254L), two of the zinc coordinating residues in the β-CASP domain of Artemis, cause RS-SCID in humans [34,44,55]. Patients with these inherited mutations suffer from impaired V(D)J recombination, leading to underdeveloped B and T lymphocytes. The importance of histidine 254, has been highlighted by de Villartay *et al.* [44], who showed that the full-length H254A Artemis variant is unable to carry out V(D)J recombination *in vivo* and has no discernible endonucleolytic activity *in vitro*.

### Comparison of Artemis structure with 6WO0 and 6WNL

During the preparation of this manuscript a structural study on the catalytic core of Artemis was published [56]. This study described reported two Artemis structures (PDB: 6WO0 and 6WNL) that are similar to our Artemis structure (PDB code 7AF1) (backbone RMSDs of 0.48 Å and 0.54 Å respectively), with identical relative positioning of the MBL and β-CASP domains (Figure 4A and Suppl. Figure 4A). The only significant difference was that whilst we refined our structure with two zinc ions in the active site, both of the crystal forms reported by Karim *et al.* were modelled with a single active site zinc ion (Zn1), reinforcing the proposal of weaker metal ion binding at the Zn2 site.

**Figure 4:**
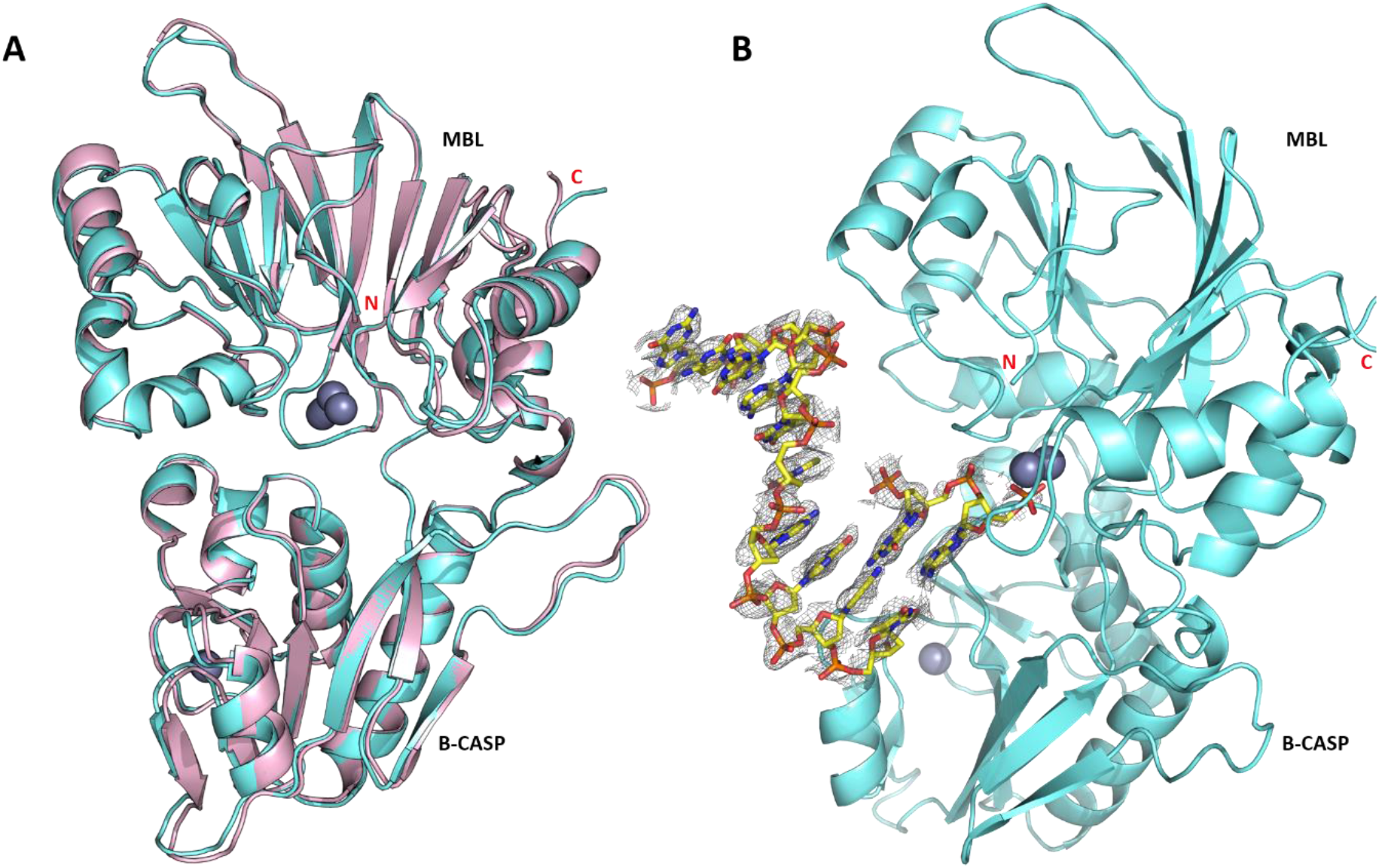
Overall structure representation of the Artemis /SNM1C fold. **A**. Overlay of our wt Artemis structure (PDB code: 7AF1) (pink) with that of Karim *et. al. PDB code:* 6WO0 (aquamarine) (backbone RMSD 0.48 Å). **B.** Re-analysis of the latter (6WO0) structure re-refined with a DNA molecule present (PDB code; 7ABS). The 2Fo-Fc map (contoured at 0.6σ in pymol) is represented by grey mesh surrounding the DNA (yellow).

### Re-analysis of the 6WO0 and 6WNL structures

An unusual aspect of the Karim *et al.* structures is that both crystal forms were obtained in the presence of DNA and were reported to require DNA for their growth; the crystals showed a fluorescence signal supporting the presence of DNA (the oligonucleotides used contained a cyanine dye fluorophore), yet neither of the models presented contain DNA. The authors referred to some broken stacking electron density in 6WNL in a solvent channel and a patch of unsolved density approaching the active site in 6WO0, but state that the DNA “did not bind to the protein in a physiological way, and likely bound promiscuously to promote crystallization” [56].

We performed a careful re-examination of these structures looking closely at the residual electron density. For the 6WNL structure we were able to locate a distorted duplex DNA of around 13 base pairs which we propose may be the product of duplex annealing of the oligonucleotide used for crystallization (a semi-palindromic 13-mer that was designed to form a hairpin with phospho-thioate linkages in the single-stranded region) (Suppl. Figure 4B). For this structure, we are in general agreement with Karim *et al* that the DNA does not appear to make meaningful interactions with the protein that inform on the mechanism of nuclease activity, although this mode of association with DNA may possibly be relevant to alternative binding modes relating to higher order complexes containing Artemis. By contrast, for the 6WO0 structure we were able to confidently build a DNA molecule that contacts the Artemis active site in a manner that we believe to be relevant to the Artemis nuclease activity.

Our model contains an 8-nucleotide 5’-single-stranded extension with a short 2-base pair region of duplex DNA that reaches into the Artemis active site making close contacts with the metal ion centre in a manner consistent with the proposed catalytic mechanism (Figure 4B). The sequence of the longest strand corresponds to the 10-nucleotide cy-5 labelled strand (cy5-GCGATCAGCT) with some residual density at the 5’-end that may be attributed to the cyanine fluorophore which we did not include in our model. The complementary strand used for crystallization was 13-nucleotides long and was intended to produce a 5’-overhang, but only two bases and three phosphates could be located in the density. The abrupt manner in which the electron density apparently disappears from either end of this strand suggests that this is the product of a cleavage reaction, although it is possible that remaining nucleotides are not located due to disorder.

The analysis of electron density at this site is complicated by the proximity to a crystallographic 2-fold symmetry axis, which brings a symmetry copy of the DNA molecule into a position where atoms partially overlap and the extended 5’ strands form a pseudo duplex (Suppl. Figure 5A). The occupancy of the entire DNA molecule is thus limited to 0.5, and the lower occupancy is reflected in the electron density map which requires a lower contour level than would usually be applied (Suppl. Figure 5B). After carefully building and refining the afore-described DNA bound model, significant positive electron density was revealed for the second metal ion (Zn2 site) which we also included in the model with the same occupancy (0.5) as the DNA. Our model was refined to similar crystallographic R-factors as 6WO0 and has been deposited with PDB accession number 7ABS (refinement statistics are given in Table I).

### Model for Artemis DNA binding

Using the crystallographically observed DNA as a template we were able make a model for Artemis binding to a longer section of double-stranded DNA by complementing unpaired bases on the single-stranded DNA overhang with canonical base pairs, whilst maintaining acceptable geometry of the sugar phosphate backbone (Figure 5A). The duplex section of this model deviates slightly from the ideal B-form geometry [57], in a manner that is reminiscent of certain transcription factor DNA complexes [58,59]. We have also extended the metal ion contacting strand by three nucleotides to form a 5’-overhang; the positioning of the overhang nucleotides is more speculative, nevertheless it was possible to avoid clashes with protein residues whist maintaining relaxed geometry (Figure 6C).

**Figure 5:**
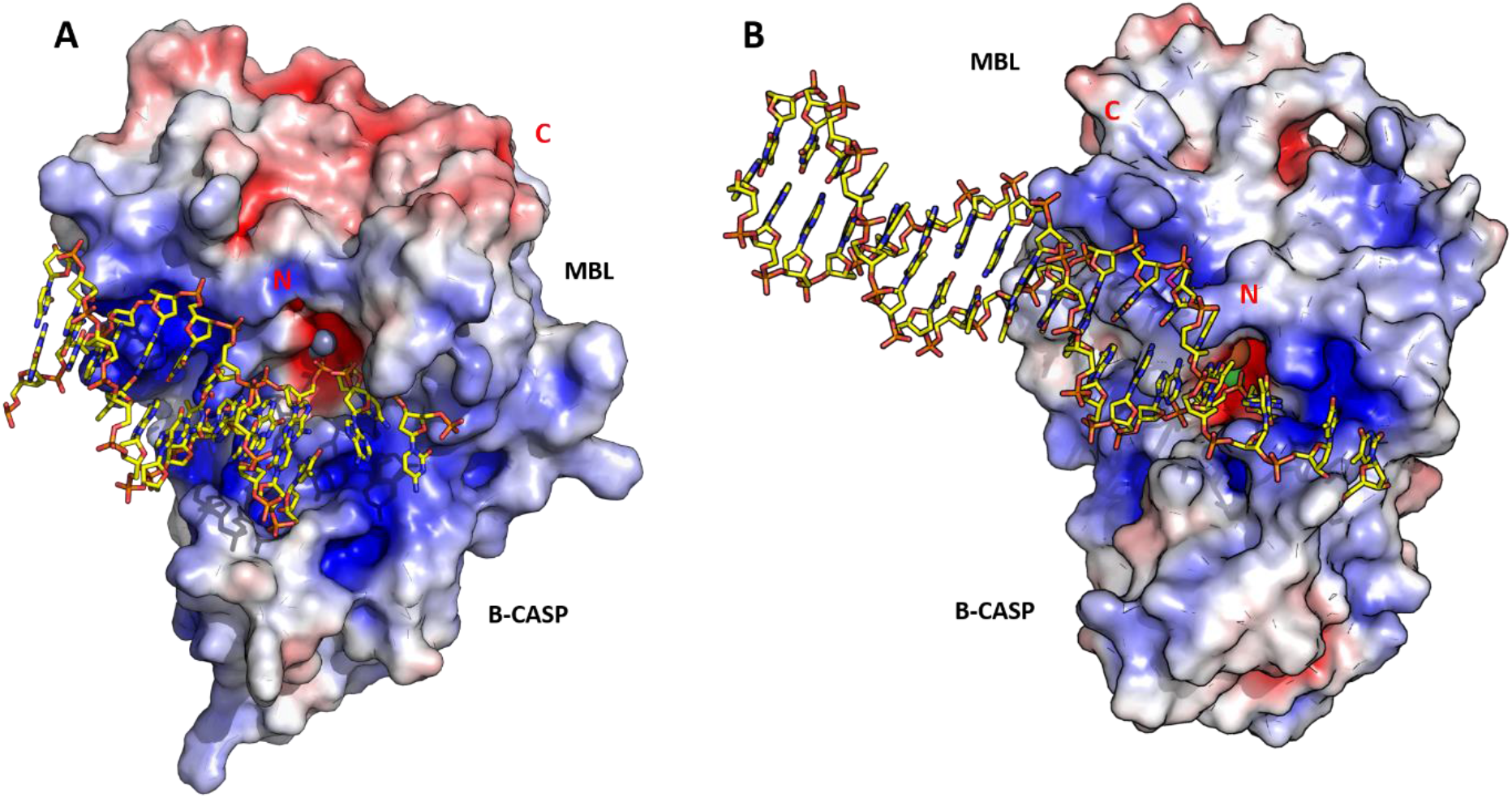
Electrostatic surface potentials of DNA bound model for Artemis/SNM1C (A) and Apollo/SNM1B (B). The blue colour represents a more electropositive surface potential and the red show a more electronegative cluster. The active site contains the two metal ions represented in grey sphere for Zinc, orange sphere for Iron and green sphere for Nickel ion. N- and C-terminal of the protein are indicated in red. The electrostatic surface potentials were generated using PyMOL (electrostatic range at +/− 5).

**Figure 6:**
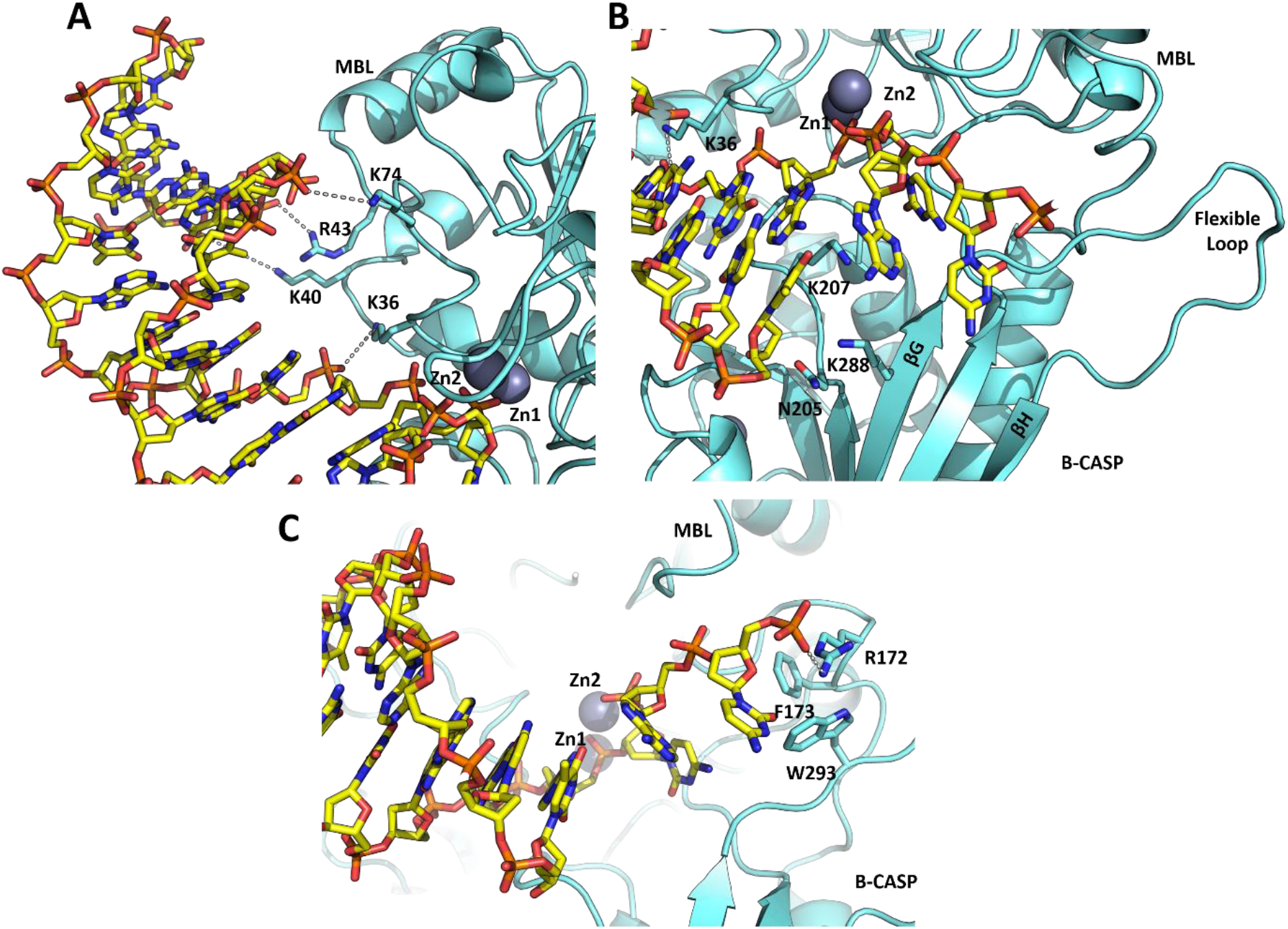
Proposed interactions of DNA with Artemis (PDB: 7ABS). Model for DNA binding to Artemis showing the residues contacting DNA. The two zinc ions at the active site are represented by grey spheres. **A.** A row of positively charged residue is on the surface of the MBL domain interact with the phosphate backbone of the DNA. **B.** A DNA overhang is located at the active site. A cluster of polar residues (N205, K207 and K288) is located in the β-CASP domain. The extended flexible loop, which is unique to Artemis compared to SNM1A and SNM1B, that connects b27 and b28 is indicated on the right. **C.** The DNA overhang forms a hydrogen bond with Arginine 172 and interacts with a cluster of hydrophobic residues at the interface between MBL and β-CASP domains.

In the extended DNA complex model, Artemis contacts both strands of the DNA model in several areas; notably a single phosphate lies above the di-metal ion bearing active site and ligates to both metal ions in the same manner as observed in structures of related enzymes with phosphate or phosphate-containing compounds (Baddock *et.al.* 2020) [60]. The two downstream nucleotides on this strand pass close to the protein surface, forming possible interactions with both the main chain of Asp37 and sidechain of Lys36, whilst subsequent nucleotides are not close to the protein (Figure 6A). The overhang portion of this strand continues with a slightly altered trajectory, potentially contacting Artemis in the vicinity of the cleft separating the MBL and β-CASP domains, with the potential to form favourable interactions with both positively charged (Arg 172) and aromatic residues (Phe 173 and Trp 293) (Figure 6C).

The complementary strand forms interactions with the protein *via* backbone contacts that span a 4-nucleotide stretch between 5- and 8-bases from the 3’-terminus that contact positively charged sidechains in the MBL domain (Lys36, Lys40, Arg43, and Lys74) (Figure 6A). The 3’-end of this strand apparently terminates directly above a cluster of polar or positively charged residues in the β-CASP domain (Lys207, Lys288, Asn205) (Figure 6B). Whilst the experimental (as used in co-crystallisation) DNA substrate and our model both contain a 3’-hydroxyl group, the model implies that addition of a 3’-phosphate could be accommodated and may be expected to make favourable interactions with the basic cluster of residues. Thus, our model illustrates a preferred binding mode for Artemis for DNA with a 5’-overhang binding at the junction between double- and single-stranded regions, and the expected product of this reaction would be a blunt ended DNA with a 5’-phosphate. In the case of hairpin DNA substrates our model indicates the possibility for Artemis to accommodate a loop connecting the two strands possibly of around 4-nucleotides or more, with the cleavage product being DNA with a 3’-overhang cleaved from the last paired base of the duplex.

### Comparison of the Artemis DNA binding mode with that of other nucleases

We have recently determined the structure of SNM1B/Apollo in complex with two deoxyadenosine monophosphate nucleotides and through a similar process of extrapolation to that outlined above we have independently built a model for SNM1B binding to DNA containing a 3’-overhang (one of its preferred substrates) (Baddock *et.al.* accompanying paper). The overall mode of DNA binding is similar in the two models (Figure 5), with the two DNA duplexes being roughly parallel and forming contacts to similar regions on the MBL domain. The most important differences lie in the nature of the contacts formed to the active site and the paths of the various overhangs. In the SNM1B model extensive contacts are made to the 5’-phosphate in a well-defined phosphate binding pocket. Both human SNM1A and SNM1B are exclusively 5’-phosphate exonucleases, with most of these phosphate binding residues being highly conserved (sequence alignment Suppl. Figure 1 in yellow) (Baddock *et. al* accompanying paper). Interestingly, Artemis lacks these key phosphate binding residues and the 5’-phosphate binding pocket of SNM1A and SNM1B. Instead, this pocket in Artemis is partially filled by the side chain of Phe318. These contacts appear to define a high-affinity binding pocket exclusively for the 5’-phosphate of the DNA terminus, thus explaining the major differences in nuclease activities within the family, i.e., SNM1A and SNM1B being exonucleases and Artemis being an endonuclease. Further differences between Artemis and SNM1B/SNM1A are found in the loop connecting β-strands G and H (using Artemis numbering), which in Artemis is significantly longer and occupies a different position contacting residues in the MBL domain (Figure 6B), compared to the loops in SNM1B and SNM1A that form part of the phosphate binding pocket and make potential contacts with the 3’-overhang. This loop displacement in Artemis may contribute to its ability in accommodating DNA substrates with either 5’-overhangs or hairpins, thus facilitating its function as a structure specific endonuclease.

Interestingly, the surface of the Artemis interface between the MBL and β-CASP domains contains a belt of positively charged residues (Figure 5A). These positive surface charges are proposed to facilitate productive DNA binding to the active site, consistent with our DNA binding model. One of the striking differences, at least in the available structures, is that the active site of Artemis is more open compared to those of SNM1A or SNM1B. This openness may reflect an ability to accommodate different substrate conformation including hairpins, 3’- and 5’-overhangs, as well as DNA flaps and gaps. Both human SNM1A and SNM1B appear to have a more sequestered active site that would only fit a single strand of DNA, which is consistent with previous findings on their preferred substrate selectivity [10].

### Biochemical characterisation of truncated Artemis catalytic domain (aa 1-361)

To investigate the activity of our different versions of recombinant Artemis, we performed nuclease assays using radiolabelled DNA substrates. We compared the catalytic domain purified using IMAC (which contained Ni^2+^ in the active site) with protein purified using ion exchange (and avoiding IMAC), which contained predominantly Zn^2+^. We also tested the activity of full-length phosphorylated Artemis. The results show that both truncated enzymes have identical activities, which is also very similar to that of the full-length enzyme (Suppl. Figure 6).

One notable difference between our full-length protein and that reported by Ma *et. al* [18], is that our full-length protein is active in the absence of DNA-PKcs. Intact protein mass spectrometric analysis of our full-length protein shows that the protein has undergone up to five phosphorylation events (Suppl. Figure 7). Poinsignon *et al*. have shown that Artemis is constitutively phosphorylated in cultured mammalian cells and is the target of additional phosphorylation in response to induced DNA damage [61]; it is interesting that the capacity to phosphorylate Artemis to produce an active form is also conserved in insect cells. We observed exonuclease activity with full-length Artemis at 10 nM (Suppl. Figure 8), though this was weak compared to its endonuclease activity at the same concentration. We observed no exonuclease activity for the truncated Artemis construct, it is possible that phosphorylation alters the balance between endonuclease and exonuclease activity, though the biological relevance of this, if any, remains to be validated.

As mentioned above, both human SNM1A and SNM1B require a 5’-phosphate for their activity [3,9,15]. To investigate whether there is a similar requirement for Artemis, we tested the activity of truncated Artemis against single-stranded and overhang DNA substrates with different 5’-end groups, including a phosphate, hydroxyl group, and biotin groups (Figure 7A). The results imply that, at least under the tested conditions, Artemis is agnostic to the different end modifications, exhibiting comparable digestion of all substrates.

**Figure 7:**
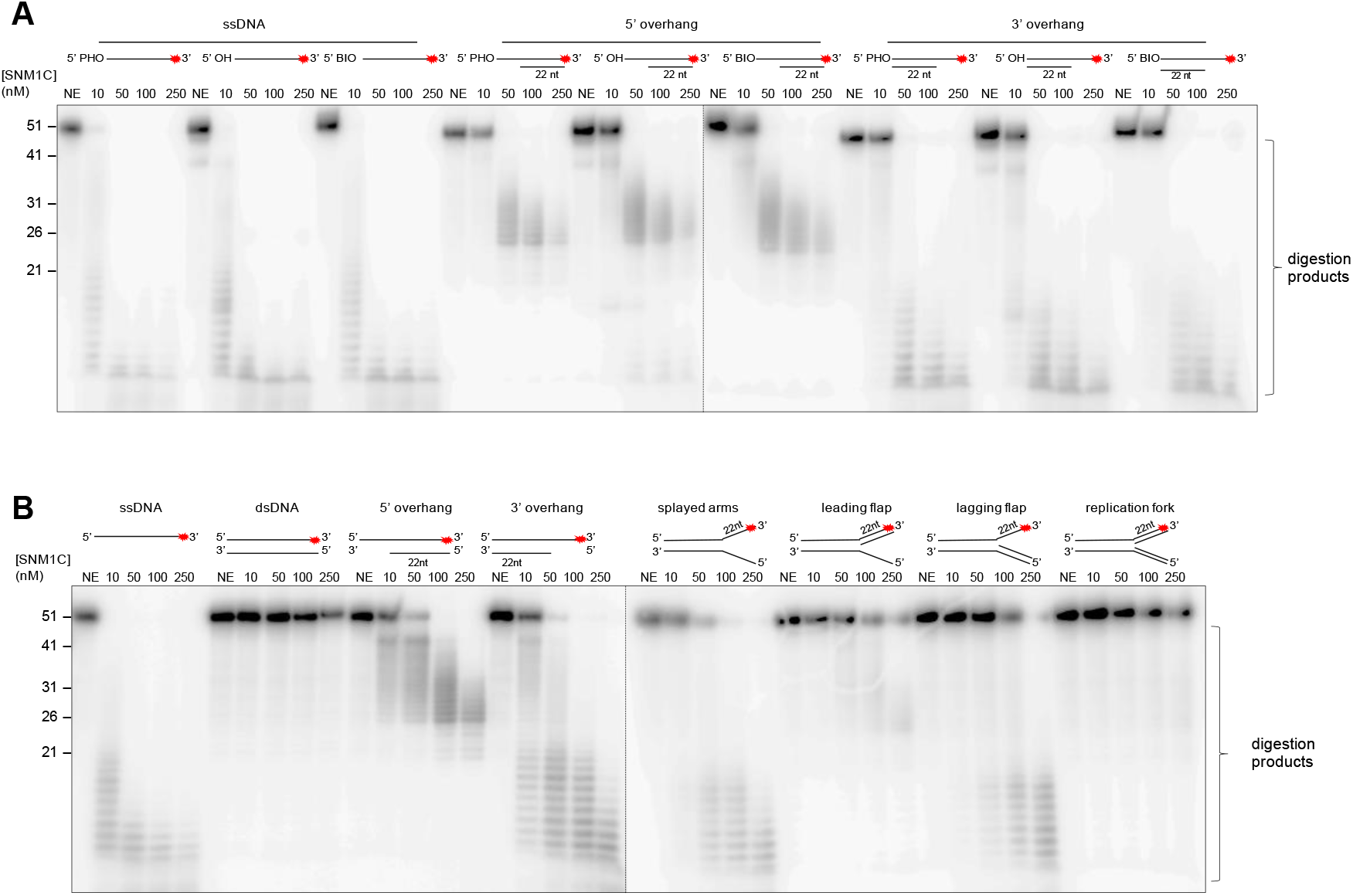
Nuclease assay utilising truncated Artemis (aa 3–361) with various DNA substrates. **A.** The nuclease activity of Artemis is indifferent to the 5’ end group, indicative of true endonuclease activity. Increasing concentrations of Artemis from 0 (NE; no enzyme) to 250 nM incubated with 10 nM ssDNA with either a 5’ phosphate, 5’ hydroxyl, or 5’ biotin moiety for 45 min at 37°C. **B**. Artemis is able to cleave DNA substrates containing single-stranded regions. Increasing amounts of Artemis incubated with structurally diverse DNA substrates (10 nM) for 45 min at 37°C. Products for A and B were analysed by 20% denaturing PAGE. The DNA substrates utilised are represented at the top of the lanes and a red asterisk indicates the position of the 3’ radiolabel. The positions of DNA size markers run as a reference are indicated on the left, with sizes in nt.

Extensive evidence demonstrates that full-length Artemis in complex with DNA-PKcs has structure specific endonuclease activity [17,18,27]. These studies reported that Artemis can digest substrates including overhangs, hairpins, stem-loops, and splayed arms (pseudo-Y). To investigate the activity of truncated Artemis catalytic domain (aa 1–361) we performed nuclease assays using a variety of radio-labelled DNA substrates (Figure 7B). The results show that truncated Artemis has substrate specific endonuclease activity, with a preference for single-stranded DNA susbstrates, and those that contain single stranded character (e.g. 5’- and 3’-overhangs, splayed arms, and a lagging flap structure), compared with double stranded DNA structures (e.g. ds DNA and a replication fork). This is in accordance with previous research, where Artemis has been reported to cleave around ss-to dsDNA junctions in DNA substrates (perhaps cite Chang et al, 2015 for this). The truncated Artemis catalytic domain also exhibits hairpin opening activity, in accordance with what has previously been reported (Suppl. Fig. 9). On a duplex substrate (YM117 from Ma *et al*) [18] with a 20 nt hairpin region, Artemis cleaves adjacent to the hairpin, consistent with previous data. It is clear that truncated form of Artemis exhibits nuclease activity closely comparable to the phosphorylated full-length Artemis protein [18], indicating that the structural studies presented here reveal mechanistic insights of direct relevance to the DNA-PKcs-associated form of Artemis that engages in end-processing reactions *in vivo*.

### Structural and biochemical characterisation of Artemis point mutations

Previous site-directed mutagenesis studies by Pannicke *et al.* targeting the metal ion coordinating residues in the active site (D17N, H33A, H35A, D37N) of full-length Artemis (aa 1–692) established the importance of active site motifs 1–4 for activity [27]. Each of these substitutions markedly reduced or abolished Artemis’ ability to carry out its role in V(D)J recombination *in vitro*.

We mutated, expressed, purified, and crystallised three forms of truncated Artemis (aa 1–361) with substitutions in several of these metal ion co-ordinating residues, i.e. D37A, H33A, and the Omenn Syndrome patient mutation H35D [31,55]. We found that the overall architecture of the three variants is almost identical to the WT (Figure 8A). The D37A structure retains one Zn ion (Zn1) whilst losing the second, (Zn2) (Figure 8B) in the active site. Both the H33A and H35D variants additionally exhibited loss of the Zn1 ion. All three variants retained the Zn ion in the zinc finger-like motif of the β-CASP domain.

**Figure 8:**
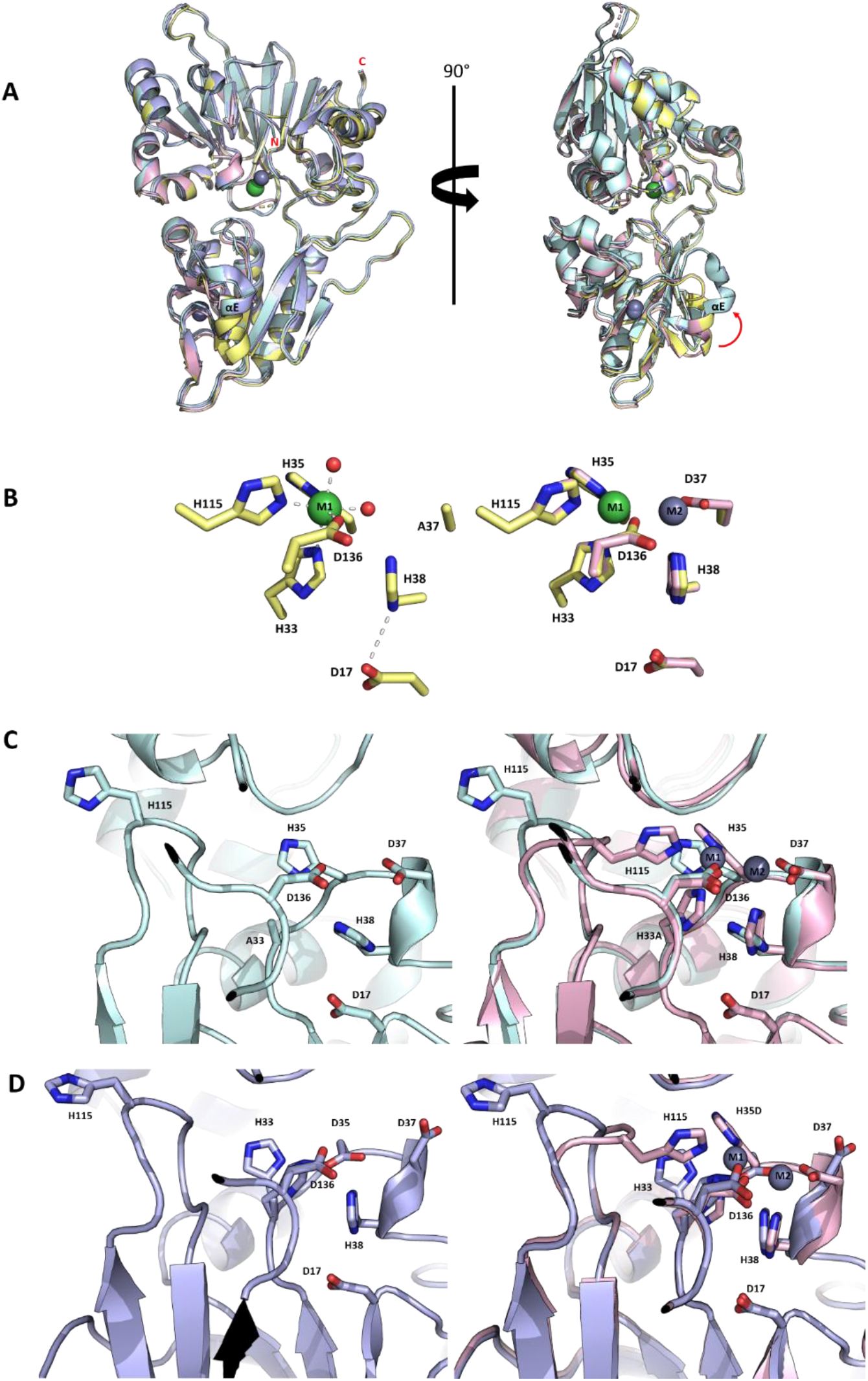
Views from structure of Artemis D37A, H33A and H35D variants. **A.** An overlay of the four Artemis structures: WT (PDB:7AF1) in pink, D37A variant (PDB:7AFS) in yellow, H33A variant (PDB:7AFU) in cyan and H35D patient mutation (PDB:7AGI) in blue, showing the general architecture of the three variants are the same as the W structure. The nickel ion is represented as green sphere, and the zinc ions as grey spheres. The movement of helix αE in the β-CASP domain is indicated by a red arrow. **B.** Left: The active site of D37A variant has a single nickel ion with two complexing water molecules (red spheres); Right: the active site residues of WT Artemis (pink), superimposed with those of the D37A variant (yellow). Aside from loss of the second metal in the D37A mutant, there is little movement at the active site. **C.** Active site residues of the H33A variant (cyan; left) and an overlay (right) with WT Artemis (pink). The two distinguishing features of H33A variant are a lack of metal ions and movement of the loop containing His115. **D.** The active site of the H35D variant (blue; left) and an overlay (right) with WT Artemis (pink). The H35D point substitution is present in patients with Omen syndrome. This variant lacks both metal ions, similarly to the H33A variant.

The position of the active site residues and the surrounding residues in the D37A variant, superposed perfectly with WT Artemis. As previously mentioned, Asp37 can adopt two conformations as seen in 6TT5 structure, noting that the coordinated zinc ion is generally present at about 30% occupancy in both of the WT Artemis structures (PDB 6TT5 and 7AF1). Therefore, it is unsurprising that mutation of Asp37 to alanine results in the loss of Zn2. Differential scanning fluorimetry (DSF) experiments were carried out to investigate the stability of the three variants. DSF analysis showed that the D37A variant has similar thermal stability as the WT, suggesting that the protein is stable and folded in the presence of a single metal ion in the active site, whilst both H33A and H35D variants were substantially destabilized with ΔT_m_ around −13 °C compared to the WT Artemis (Suppl. Figure 10).

Histidines 33 and 35 are the first two histidine residues in the HxHxDH motif (motif 2) in the MBL domain. Their role is to coordinate the first metal ion (Zn1 site) in the catalytic site. In the absence of metal ions in the catalytic site, the loop comprising residues 113–119 moves away from the active site (Figure 8C and D). Another small rearrangement occurs in helix α8 (residues 348–358) of the MBL domain. In both the H33A and H35D variants, helix α8 moves slightly closer toward strand β14, compared to the WT and D37A variant. Surprisingly, the biggest rearrangement occurs in β-strand E (residues 268–270) and α-helix E (residues 261–267); both located near the zinc finger motif in the β-CASP domain (Figure 8A). In H33A and H35D variants, both β-strand E and α-helix E shifted upward and away from the zinc finger like motif. These conformational changes may suggest some allosteric regulation in terms of substrate binding and catalytic activity of the enzyme.

We also tested the activity of the D37A, H33A, and H35D Artemis variants *in vitro* using single-stranded 3’ end radiolabelled DNA as a substrate in a gel-based assay. All three variants lost their ability to digest the DNA substrate *in vitro* (Figure 9). These observations are in agreement with the results obtained with full-length variants by Ege *et al. and* Pannicke *et al.* [27,31]. Their studies show that the full-length Artemis variants H33A, H35D, and D37N are able to interact with and be phosphorylated by DNA-PKcs, however, have lost the ability to digest DNA substrates *in vitro*. The combined results reveal the importance of the HxHxDH motif and highlight the importance of the di-metal catalytic core in the SNM1 family, not only in directly catalysing hydrolysis, but also likely in conformational changes involved in catalysis [43].

**Figure 9:**
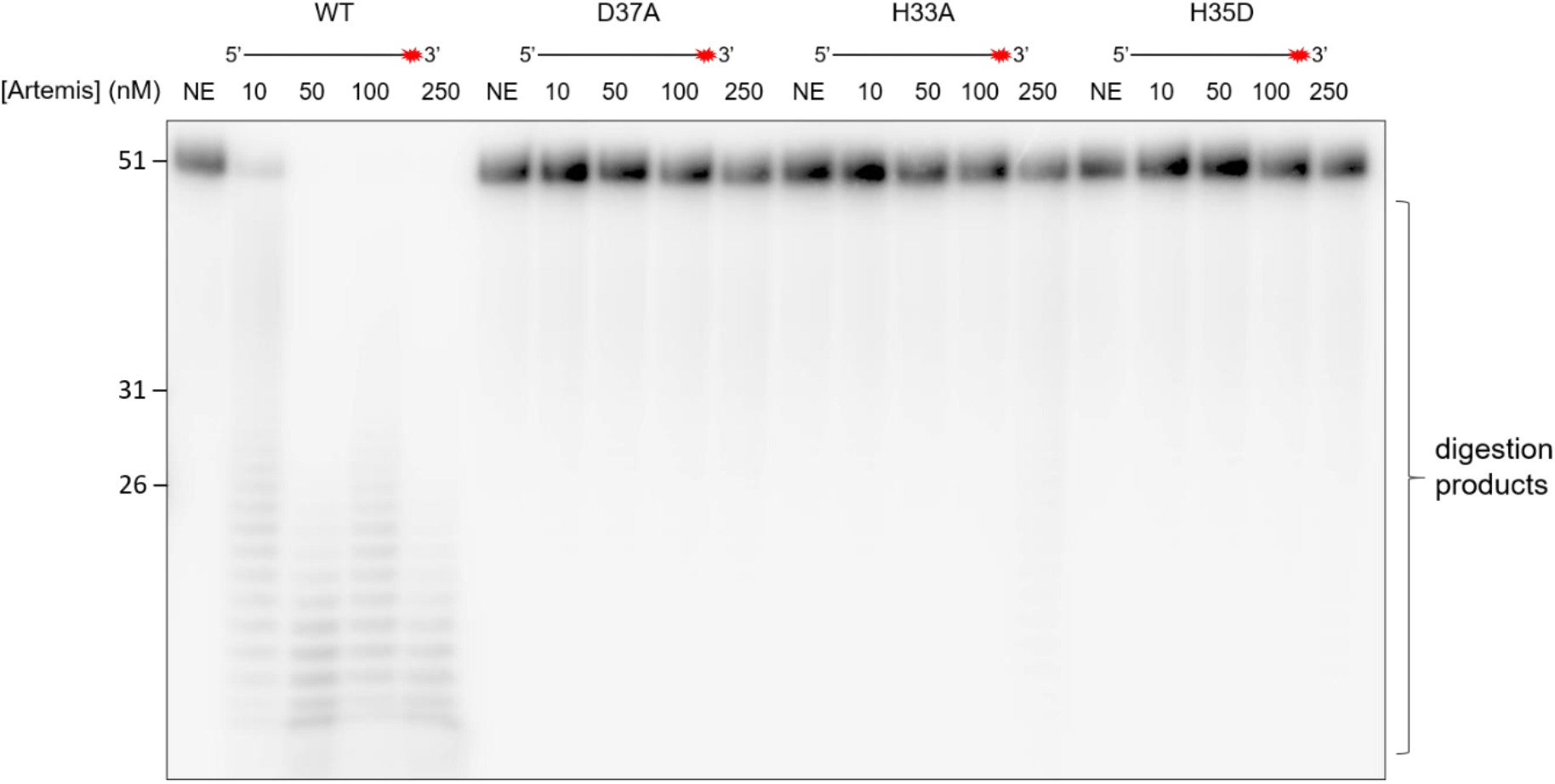
Comparing the activity of Artemis variants vs WT protein. Increasing amounts (from 0 to 250 nM) of WT and mutant Artemis proteins (as indicated) were incubated with 10 nM of 51 nucleotide ssDNA substrate for 30 min at 37 °C. Reaction products were subsequently analysed by 20% denaturing PAGE. The size (in nucleotides) of the marker oligonucleotides are indicated on the left-hand side of the corresponding bands.

### Identification of small molecule inhibitors of Artemis

Radiotherapy is a mainstay of cancer therapy; its effectiveness relies on inducing DNA double-strand breaks (DSBs) that contain complex, chemically modified ends that must be processed prior to repair [62,63]. The canonical non-homologous end-joining (c-NHEJ) pathway repairs ~80% of DNA double-strand breaks in mammalian cells [20,23]. Therefore, combining radiation therapy in conjunction with c-NHEJ inhibitors could selectively radiosensitise tumours. Weterings *et al*. have reported a compound that interferes with the binding of Ku70/80 to DNA, thereby increasing sensitivity to ionising radiation in human cell lines [64]; ATM inhibitors are also in advanced clinical development and represent the most developed strategy to inhibit DSB repair to increase the efficacy of radiotherapy [65]

Artemis, along with SNM1A and SNM1B, possess a conserved MBL-fold domain that is similar to the true bacterial MBLs. Previous studies on human SNM1A and SNM1B/Apollo, showed that ceftriaxone (Rocephin), a widely used β-lactam antibacterial (third generation cephalosporin) inhibits the nuclease activity of both SNM1A and SNM1B [66]. To investigate if this class of β-lactam anti-bacterial compounds could inhibit Artemis we performed fluorescence-based nuclease assays with three cephalosporins, i.e. ceftriaxone, cefotaxime and 7-aminocephalosporanic acid (Figure 10A). The results show that neither cefotaxime nor the parent compound, 7-aminocephalosporanic acid, potently inhibit Artemis’ activity, whilst ceftriaxone inhibits Artemis with a modest IC_50_ of 65 μM (Figure 10B).

**Figure 10:**
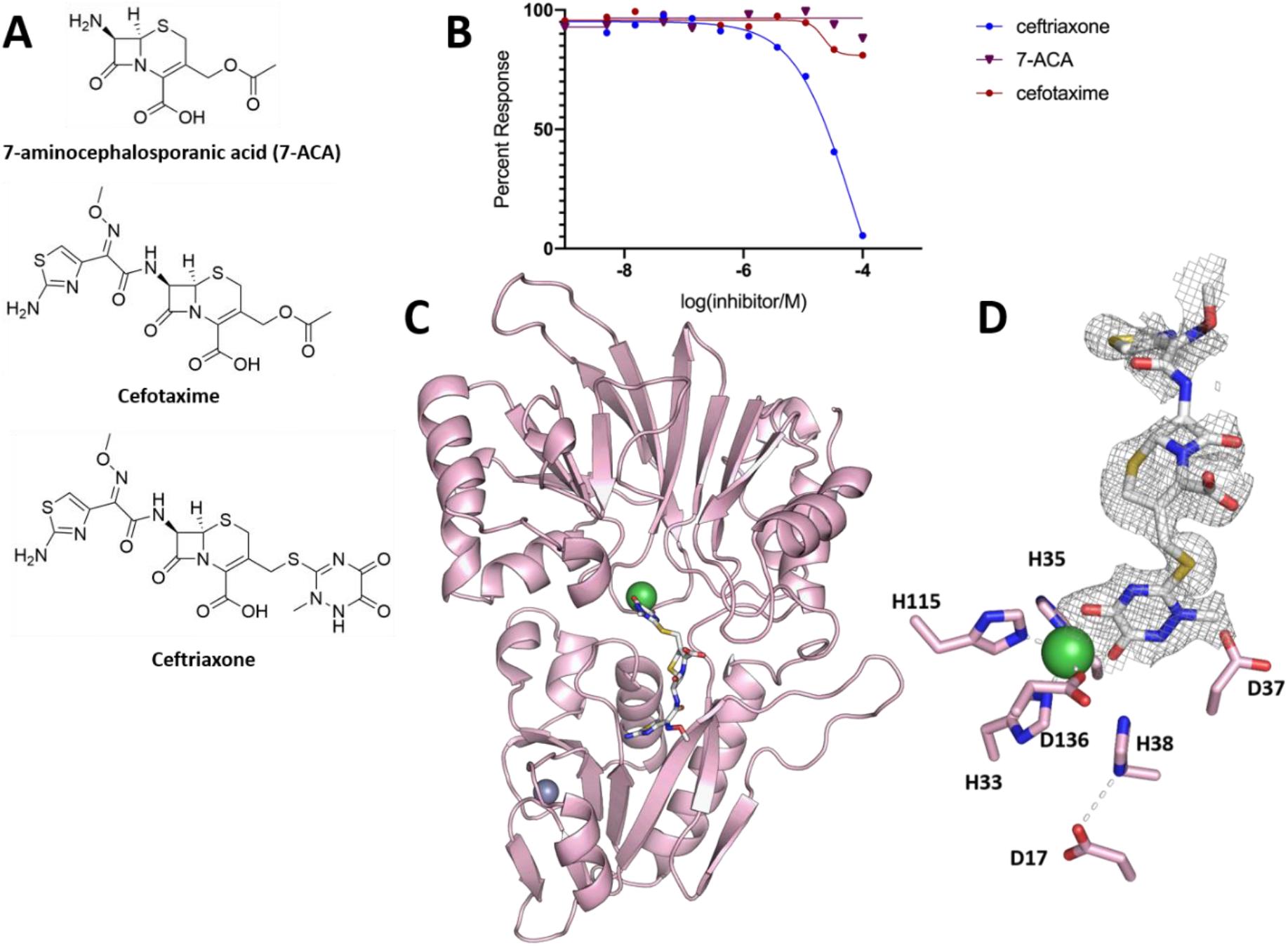
Artemis inhibition by β-lactam antibiotics. **A.** Structures of selected β-lactam antibacterial. **B.** Inhibitor profiles of β-lactams on the nuclease activity of Artemis was assessed *via* a real-time fluorescence-based nuclease assay. **C.** Cartoon representation of the structure of truncated Artemis (aa1-361) with a ceftriaxone molecule (in white) bound at the active site. **D.** Active site residues with the electron density (Fo-Fc) contoured map around the modelled Ceftriaxone. The map is contoured at the 1.0 σ level and was calculated before the Ceftriaxone molecule was included in the refinement.

We solved the structure of ceftriaxone bound to the catalytic domain of Artemis (purified by IMAC) at 1.9 Å resolution (Figure 10C) by soaking an Artemis crystal with ceftriaxone. This structure was solved by molecular replacement (using PDB: 6TT5 as a model), in the space group P1 with one protein molecule in the asymmetric unit. As before, in this structure Artemis possesses the canonical bilobar MBL and β-CASP fold with an active site containing one nickel ion, possibly related to the purification method.

Ceftriaxone binds to the protein surface in an extended manner making interactions with the active site, towards the β-CASP domain (Figure 10 C). There is no evidence for cleavage of the β-lactam ring nor of loss of the C-3’ cephalosporin side chain, reactions that can occur during ‘true’ MBL catalysed cephalosporin hydrolysis. The electron density at the active site clearly reveals the presence of the ceftriaxone side chain in a position to coordinate the nickel ion (at the Zn1 site) replacing water molecules (waters 72 and 106) compared with the apo structure (Figure 10D). Despite the conservation of key elements of the active site of the MBL fold nucleases and the ‘true’ b-lactam hydrolysing MBLs [67], ceftriaxone, does not interact with the nickel ion *via* its b-lactam ring (as occurs for the true MBLs), but *via* both carbonyl oxygens of the cyclic 1, 2 diamide in its sidechain (Ni-O distances: 2 Å and 2.2 Å), i.e. it is not positioned for productive b-lactam hydrolysis. The amino-thiazole group (N7) of ceftriaxone forms hydrogen bonds with the side chain of Asn205, while the S1 of the 7-aminocephalosporanic acid core of the compound interacts with the hydroxyl of Tyr212 through an ethylene glycol molecule. The rest of the molecule appears to be flexible. The binding mode of ceftriaxone to Artemis shown in Figure 10C is near identical to that observed for ceftriaxone with SNM1A (PDB: 5NZW) structure (Suppl. Figure 11).

One notable difference between the Ceftriaxone-bound Artemis structure and the apo structure, is the loss of a second metal ion at the active site (Figure 10D). In the apo structure (Figure 2C), this zinc ion is coordinated by residues Asp37, His38, and Asp130. With a single metal coordination in the ceftriaxone bound structure, the Asp37 side chain is positioned away from the active site (Figure 10D), as seen in the nickel bound (PDB: 6TT5) structure (Figure 2E).

To investigate the possibility of inhibiting Artemis through binding to the zinc finger motif in the β-CASP domain, we used the fluorescence-based nuclease assay to test three compounds known to react with thiol groups present in zinc fingers and which result in zinc ion displacement, i.e. ebselen [68], auranofin [69] and disulfiram [70]. We found that both ebselen and disulfiram inhibit Artemis with IC_50_ values around 8.5 μM and 10.8 μM respectively, whilst Auranofin inhibits less potently (IC_50_ 46 μM) (Suppl. Figure 12), indicating additional possible inhibitory strategies.

## DISCUSSION

The DCLRE1C/Artemis gene was first discovered in 2000, following work with children with severe combined immunodeficiency disease (SCID) [71]. Subsequent studies have shown that Artemis is a key enzyme in V(D)J recombination [16,17,72] and the c-NHEJ DNA repair pathway [21,44,73]; and that it is structure specific endonuclease, and member of MBL fold structural superfamily [2]. Our structures of wild-type and catalytic site mutants of SNM1C/DCLRE1C or Artemis protein show that, like SNM1A and SNM1B/Apollo and the RNA processing enzyme CPSF73, Artemis has a typical α/β-β/α sandwich fold in its MBL domain and has a β-CASP domain, the latter a characteristic feature of MBL fold nucleases. However, both our Artemis structures and those recently reported by Karim *et. al* [56] reveal a unique structural feature of Artemis in its β-CASP domain that is not reported in other human MBL enzymes, i.e. a classical zinc-finger like motif. Moreover, collectively, these structures allow us to assign a likely mode of DNA substrate interaction for Artemis.

The role of the newly-described zinc-finger like motif remains unknown. However, zinc-finger motifs are common structural features in DNA binding proteins such as transcription factors [50,54], but are also observed in a substantial number of required and accessory NHEJ proteins [52]. These zinc fingers provide structural stability and enhance substrate selectivity rather than being involved in catalytic reactions, and we propose that this is likely to be the case for Artemis. The fact that the residues (His 228, His 254, Cys 256, and Cys 272) that are involved in the zinc-finger like motif are highly conserved across different Artemis species suggests the importance of this structural feature. Furthermore, point mutations in His 228 and His 254 (H228N and H254L) have been reported in patients with a SCID phenotype [55].

The presence of one or two metal ions coordinated by the HxHxDH motif at the active site of Artemis reflects a hallmark of the SNM1 enzyme family [9,74]; the available evidence implies that metal ion binding at one site (Zn1 site in standard MBL nomenclature) is stronger than at the other (Zn2 site). By analogy with studies on the true MBLs, these metal ions are proposed to activate a water molecule that act as the nucleophile for the phosphodiester cleavage. Our structure (PDB: 7AF1) suggests that the native metal ion(s) residing in the active site of Artemis is zinc, although a nickel ion can also occupy the same site depending on the how the protein was purified (PDB: 6TT5). Neither the presence of Ni ion in the active site, nor the truncation of the C-terminal tail appear to inhibit, at least substantially, the activity of Artemis. Thus, using radio-labelled gel-based nuclease assays, we showed that the truncated Artemis catalytic domain (aa 1–361) with either Zn or Ni ions in the active site (as observed crystallographically in the same preparations) have similar activity with the full-length Artemis construct (aa 1-693). Therefore, it seems likely that nickel ions are able to replace zinc ions in solution, but catalysis of MBL fold enzymes, including hydrolytic reactions, with metal ions other than zinc is well-precedented [67,75]

We also solved structures of three Artemis catalytic mutants; D37A, H33A, and an Omenn syndrome patient mutation, H35D. Using gel-based nuclease assays, we showed that these variants are biochemically inactive. Overall, the three variant structures are similar to the WT structure, even though H33A and H35D entirely lack any metal ions in the active site, although zinc was present in the zinc finger. Mutation of Asp37 to alanine results in the loss of the second metal in the catalytic site, likely explaining the loss of activity, although the first metal ion is still present. Note that some MBL fold hydrolases uses two metal ions (e.g.,B1 and B3 subfamilies of the true MBLs and RNase J1 from *Bacillus subtilis*) (Suppl. Figure 13A and C) [67,76] whereas others, sometimes with apparently very similar active sites, only use one metal ion (e.g. the B2 subfamily of the true MBLs and RNase J from *Staphylococcus epidermis*) [67,77](Suppl. Figure 13B and D). Thus, whilst our results support the importance of having both metals for the nuclease activity by Artemis, subtle features can influence MBL fold enzyme activity [67,74].

Following re-analysis of the Karim *et.al* structure (PDB code 6WO0), we were able to generate a model of a DNA overhang I. complex with Artemis that informs on the substrate binding mode. Our model shows that Artemis interacts with the DNA substrate in the interface between the MBL and the β-CASP domains. This interaction is mediated through the combination of polar or positive residues and aromatic residues of Artemis and the DNA substrate (Figure 6).

Artemis is the only identified MBL/β-CASP DNA processing enzyme that possesses substantive endonuclease activity. By contrast both SNM1A and SNM1B/Apollo are strictly 5’-phosphate exonucleases [3,9,15]. The recent structure of SNM1B/Apollo in complex with two deoxyadenosine monophosphate nucleotides (PDB code: 7A1F) reported by Baddock *et. al.* 2020, reveals a cluster of residues that form a 5’-phosphate binding pocket, adjacent to the metal centre. Structural sequence alignments of the three proteins shows that these residues are highly conserved in SNM1A and SNM1B (Suppl. Figure 1). Apart from Ser 317, none of these conserved phosphate binding pocket residues are present in Artemis. Instead, the pocket is partially occupied by the Phe318 side chain, which is absent in both SNM1A and SNM1B. Artemis also possesses a longer and more flexible loop connecting β-strands 27 and 28 (Figure 6B), compared to the same loop in SNM1A and SNM1B that make up part of the 5’-phosphate binding pocket. The flexibility of this loop could enable accommodation with different types of DNA structures, such as hairpins, and 5’-overhangs. These differences plausibly explain Artemis’ substrate preferences and its primary activity as a structure-selective endonuclease.

Of the three of the β-lactam anti bacterial compounds previously shown to inhibit SNM1A [66], we only observed inhibition of Artemis with ceftriaxone. Although the potency of inhibition is moderate (IC_50_ 65 μM), we were able to solve the structure of ceftriaxone in complex with Artemis. Notably, ceftriaxone does not bind with its b-lactam carbonyl located at the active site where it ligates to one zinc (or other metal) ion, but instead binds the single nickel ion in bidentate manner *via* the carbonyls of its cyclic 1,2 diamide on its C-3’ sidechain. [74,78]. Studies with the true MBLs have shown that appropriate derivatisation of weakly binding molecules can lead to highly potent and selective inhibitors.

In proof of principle attempts to inhibit Artemis though its novel structural feature compared to other MBL fold nucleases, i.e. *via* its zinc-finger like motif, we tested three covalent inhibitors with thiol-reactive groups. Ebselen, disulfiram and auranofin have the potential to interact with zinc fingers, including *via* zinc ejection with consequent protein destabilization [69,70,79,80]. Both ebselen and auranofin are reported have some antimicrobial properties [81], ebselen is in clinical trials for a variety of conditions, ranging from stroke to bipolar disorder [82], and auranofin is used for treatment of rheumatoid arthritis [83]. Recent studies have also shown that ebselen inhibits enzymes from SARS-CoV-2, i.e. the main protease (Mpro) and the exonuclease ExoN (nsp14^ExoN^-nsp10) complex [84,85]. Disulfiram is a known acetaldehyde dehydrogenase inhibitor used in treatment for alcohol abuse disorder [86]. Our results show that both ebselen and disulfiram inhibit Artemis (IC_50_s 8.5 μM and 10.8 μM, respectively), whilst auranofin is less potent (IC_50_46 μM).

Studies focussed on inhibiting the MBL fold nucleases are at an early stage compared with work on the true MBLs. The structures and assays results presented here provide starting points with established drugs, from which it might be possible to generate selective Artemis inhibitors, either binding at the active site or elsewhere (including the apparently unique zinc finger of Artemis), in order to radiosensitise cells.

## ACCESSION NUMBERS

Coordinates and structure factors have been deposited in the Protein Data Bank under accession codes 6TT5, 7AF1, 7AFS, 7AFU, 7AGI, 7APV and 7ABS.

## SUPPLEMENTARY DATA

Supplementary Data are available at NAR online.

## ACKNOWLEDGEMENT

We are very grateful to Dr Rod Chalk and Tiago Moreira for mass spectrometry, and Dr Neil Patterson for the helpful discussions and data collections at Diamond Light Source. We acknowledge Diamond Light Source for time on Beamlines I03, I04 and I24 under Proposal MX19301.

