## Supplemental Figs 1-13 and Supplemental table 1 for "Structural and mechanistic insights into the Artemis endonuclease and strategies for its inhibition"

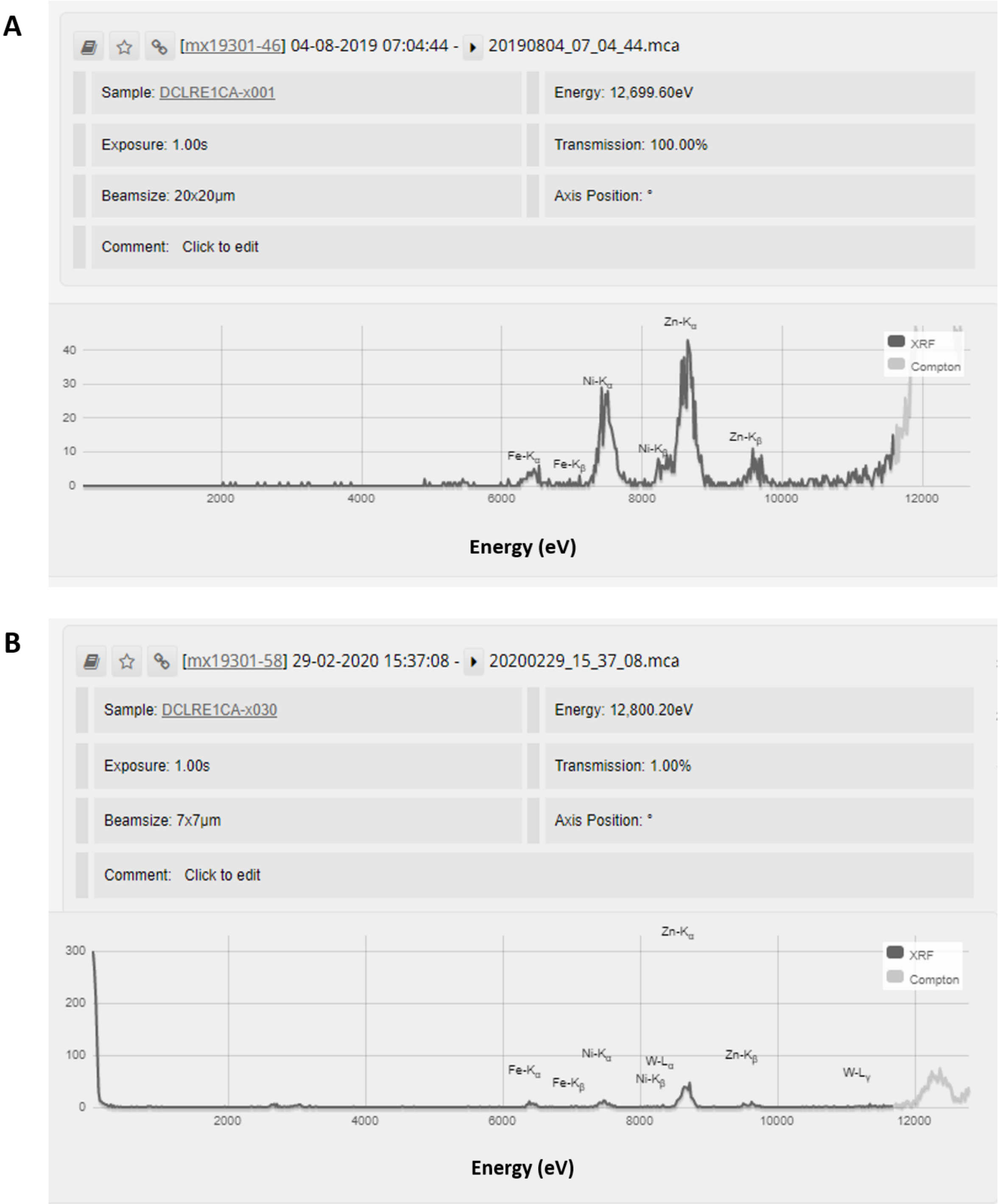


**Supplementary Figure 1: X-ray fluorescence of analysis of metal content in the WT Artemis crystals**
**A:** XRF analysis for WT Artemis crystal (PDB: 6TT5) purified using IMAC. The predominant metal species in these crystals are Ni and Zn. **B:** XRF analysis for WT Artemis crystal (PDB: 7AF1) purified without IMAC. Zinc is the predominant metal in this crystal form.

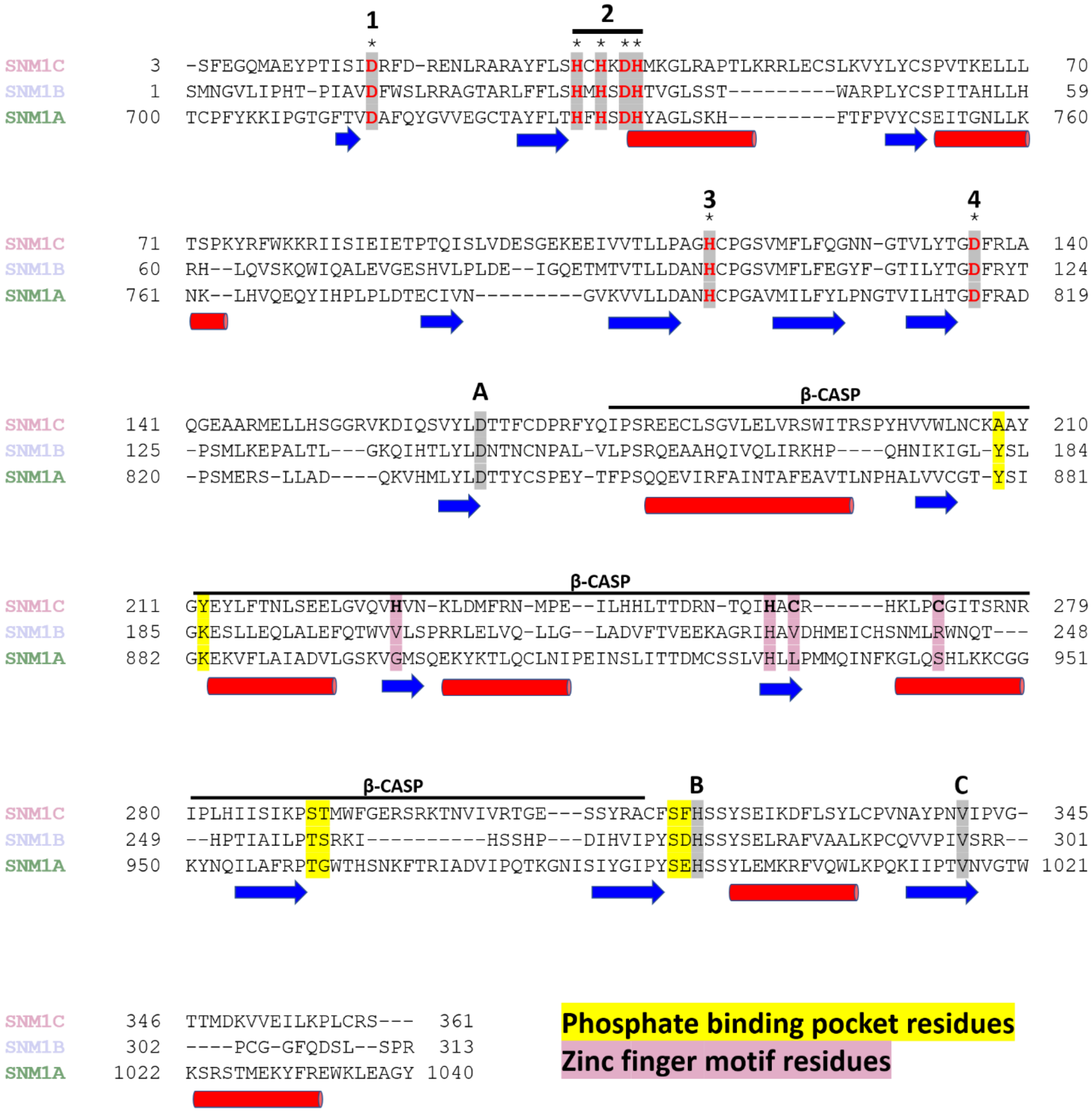


**Supplementary Figure 2: Structural sequence alignments of the human SNM1 protein family.** The structural alignment was carried out using PROMALS3D [1]. The α-helices are drawn as red cylinders and the β-strands as blue arows. The conserved MBL family motifs are labeled as 1–4 and the cannocial β-CASP motifs are labeled A–C. The conserved phoshate binding residues in SNM1A and SNM1B are highlighted in yellow. The residues that made up the novel zinc finger like structure are highlighted in pink.

```
                                                1                2
                                                *                * * **
Homo_sapiens                    MSSFEGQMAEYPTISIDRFDRENL-RARAYFLSHCHKDHMKGLRAPTLKRRLECSLKVYL    59
Rattus_norvegicus               MSSFQGQMEEYPTISIDRFDRENL-KARAYFLSHCHKDHMKGLRAPSMKRRLECSLKVFL    59
Danio_rerio                     MSSFAGRMKEYPSISLDRFDRENL-HARAYFLSHCHKDHMKGLKGPLLKRKLKFSLTVKL    59
Petromyzon_marinus              MSSFLGKTAEYPSVSIDRFDRDNV-DARAFFLSHCHQDHMVGLYSEALRKKLERSPGVHL    59
Gallus_gallus                   MSRFGGRLREYPQLSIDRFDYDNL-RARAYFLSHCHKDHMKGLRAPALRRRLQSSLKVKL    59
Python_bivittatus               MSRFGGVMREYPRLSIDRFDRENL-RARAYFLSHCHKDHMKGLRAPPMKRRLACSLKVHL    59
Xenopus_tropicalis              MSSFMGRMKEYPAISIDRFDRENL-SARAYFLSHCHKDHMKGLRAPFLKRRLQNSLKVHL    59
Chelonia_mydas                  MSSFGGRMREYPWLSIDRFDRENL-RARAYFLSHCHKDHMKGLRAPSLKRRLECSLKVHL    59
Amphimedon_queenslandica        MSVFKGSILEYPGLVVDRFNPVNCRRGRAFFLSHCHKDHMSGLDSDELLKVL-KELKIDF    59
                                ** * *   *** : :***:  *    .**:******:*** ** .  : : *  .  : :

                                                                                          3
                                                                                          *
Homo_sapiens                    YCSPVTKELLLTSPKYRFWKKRIISIEIETPTQISLVDE-ASGEKEEIVVTLLPAGHCPG    118
Rattus_norvegicus               YCSPVTKELLLTSPKYKFWENRIIAIEIETPTQVSLVDE-ASGEKEEVVVTLLPAGHCPG    118
Danio_rerio                     YCSYVTKELLLSNPRYAFWEDHIVPLELDSPTSISLIDE-STGETEDVVVTLLSAGHCPG    118
Petromyzon_marinus              YCSPVTRELLLSNWRYRFLERFTVAIEVDTATQISLMDD-KTKQKTDLVVTLLPAGHCPG    118
Gallus_gallus                   YCSPVTKELLLTNSKYAFWENHIVALEVETPTQISLVDE-TTGEKEDIEVTLLPAGHCPG    118
Python_bivittatus               YCSPVTKELLLSSPRYKFWENHIVALEVETPTQISLIEE-ASGEKEDIEVTLLPAGHCPG    118
Xenopus_tropicalis              YCSPVTKELLLTNPKYAFWENRMISIEIDTPTQISLVDE-ATGYKEDVVVTLLPAGHCPG    118
Chelonia_mydas                  YCSPVTKELLLTSPKYKFWENRIVTLEVETPTQISLLDE-ASGEKEEIVVTLLPAGHCPG    118
Amphimedon_queenslandica        YCSEVTHALLSNDPGFSHLMPYLVSVPVGETISLKLNTFRNEDQVVTVNVTLLPAGHCPG    119
                                *** **: ** ..  : .      : : :    .:.*            : **** ******

                                                 4                            A
                                                 *                                        ___
Homo_sapiens                    SVMFLFQGNNGTVLYTGDFRLAQGEAARMELLHS-GGRVKDIQSVYLDTTFCDPRFYQIP    177
Rattus_norvegicus               SVMFLFQGSNGTVLYTGDFRLAKGEVSRMELLHS-GGRVKDIQSVYLDTTFCDPRFYQIP    177
Danio_rerio                     SVMFLFEGAKGTVLYTGDFRLAVGDAARMEYLHS-GDRVKDIQSVYIDTTFFDPKYYQIP    177
Petromyzon_marinus              SVMFLFEGDHGAVLYTGDFRLPRGGVSRMGALHL-AGRIKRIKSIYLDTTFCDPMCYYIP    177
Gallus_gallus                   SVMFLFQGENGTVLYTGDFRLAKGEAARMELLHS-GTSVKDIQSVYLDTTFCDPRFYHIP    177
Python_bivittatus               SVMFLFQGENGTVLYTGDFRLARGEVARMELLHS-GSRVKDIQSVYLDTTFYDPKFYQIP    177
Xenopus_tropicalis              SVMFLFQGNSGTVLYTGDFRLAKGEVARMELLHS-GNRVKDIESVYLDTTFCDPKYYQIP    177
Chelonia_mydas                  SVMFLFQGQNGTVLYTGDFRLAKGEAARMELLHS-GTRVKDIQSVYLDTTFCDPRFYQIP    177
Amphimedon_queenslandica        SVMFLFQGDAGNVLYTGDFRLSLSDIRGCGPLHTDDGKVIEIKALYLDTTFCHPKSTNII    179
                                ******:*  * *********  .       **      :  *:::*:**** .*    *

                                                           β-CASP
                                ____________________________________________________________
Homo_sapiens                    SREECLSGVLELVRSWITRSPYHVVWLNCKAAYGYEYLFTNLSEELGVQVHVNK--LDMF    235
Rattus_norvegicus               SREECLRGVLELVRSWITRSPKHVVWLNCKAAYGYEYLFTNLSEELGVQVHVDK--LDMF    235
Danio_rerio                     SREACLAGIQQLVQDWICQSPYHVVWLNCKAAYGYEYLFTNLGQEFNSQIHVNS--LDMF    235
Petromyzon_marinus              SREDCLKGILNLVRGWITQSPQHVIWLNCKAAYGYEYLFTHLSREFGQKVHLGY--LEMF    235
Gallus_gallus                   SREECLSGILELVRSWTTLSRYHVVWLNCKAAYGYEYLFINLSEELGIKVHVNK--LDMF    235
Python_bivittatus               SREECMKGIMELVRSWVVQSPYHVVWLNCKAAYGYEYLFTNLSEELGIKVHVNK--LDMF    235
Xenopus_tropicalis              SREECLSGILELVRSWITLSPFHVVWLNCKAAYGYEYLFTNLSEEFGAKVHVNK--LDMF    235
Chelonia_mydas                  SREECLNGILELVRSWITLSRHHVVWLNCKAAYGYEYLFTNLSEELGVKVHVNK--LDMF    235
Amphimedon_queenslandica        SRDETRDIILKKVKEWLAQGPDNIVRLDCR-SFGYEHILMSLSLQLDTNIHTAGWKIRSY    238
                                **:     : : *: *   .  ::: *:*: ::***:::  *. ::. ::*      :  :

                                                           β-CASP
                                ____________________________________________________________
Homo_sapiens                    RNMPEILHHLTTD-RNTQIHACRHPKAEEYFQWSKLPCGITSRNRIPLHIISIKPSTMWF    294
Rattus_norvegicus               KNMPDILHHLTTD-RNTQIHACRHPKAEEYFQWNKLPCGMASKTKTVLHTISIKPSTMWF    294
Danio_rerio                     KKMPEILCHVTTN-RATQIHACRHPKDEEFFRANRLPCGSTAPDGIPLNIISIKPSTIWF    294
Petromyzon_marinus              QNMPDILDHVTHD-SNTQIHACRIPKDAGICEGRRLPCREKDAKGKPLHMMSIKPSTMWF    294
Gallus_gallus                   KNMPEILYHITTD-RYTQIHACRHPKDDDYVRGNRLPCGITCQNGTPLHVISIKPSTMWF    294
Python_bivittatus               KNMPEILYHVTTN-QHTQIHACRHPRDDELFRGNKLPCGITSQNGHQFHVISVKPSTMWF    294
Xenopus_tropicalis              KNMPEILSHITTD-RRTQIHACRHPVNEEFTRANRMPCGMFSDDGIPLHVISIKPSTIWF    294
Chelonia_mydas                  NNMPEILYHVTTD-RHTQIHACRHPRDDDYLRGNRLPCGMTSRNGTPLCIISVKPSTMWF    294
Amphimedon_queenslandica        SVLPQVHQCLTEDGNSTRIHACVNKKGTQSV-SGKLPCGATPTKGGTPNILTIKPSAQWF    297
                                  :*::   :* :   *:****             ::**             :::***: **

                                         β-CASP            B               C
                                ____________________
Homo_sapiens                    GERSRKTNVIVRTGESSYRACFSFHSSYSEIKDFLSYLCPVNAYPNVIPVGTTMDKVVEI    354
Rattus_norvegicus               GERTRKTNVIVRTGESSYRACFSFHSSYSEIKDFLSYICPVNAYPNVIPIGLTVDKVMDF    354
Danio_rerio                     GERTRKTSVVVKMGSSSYRACFSFHSSYLEVKDFLSYICPVNIYPNVIPLGKTVEDLTEL    354
Petromyzon_marinus              TQRSKKTKIIVRTGESSYRACFSFHSSFSEIEDFISYIKPANIFPNVIPKGKTVEDIKEM    354
Gallus_gallus                   GERIKKTNVIVRTGESTYRACFSFHSSYSEIMDFLSYIRPVNVYPNVLPVGGSEDKVMEI    354
Python_bivittatus               GERTRKTSVIMRTGMSSYRACFSFHSSYSEIQDFLSHICPENVYPNVIPVGSTEDKVKEN    354
Xenopus_tropicalis              GERTRRTNVIVRTGESSYRACFSFHSSYSEIKDFLGYIKPMNVYANVIPMGKSIEHVEKI    354
Chelonia_mydas                  GERTRKTNIVVRTGESSYRACFSFHSSYSEIKDFVGYIRPVNVYPNVIPVGSTADEVVKI    354
Amphimedon_queenslandica        LSNDKPHPSAFFPEFNLFRVLHSMHSSYSEIIEVVSYLCPVSIVPCVIPYTLGDTSLVDI    357
                                  .. :     .     . :*. .*:***: *: :.:.:: * .     *:*         : .
```

**Supplementary Figure 3: The catalytic core of DCLRE1C/SNM1C/Artemis protein sequence alignments showing conservation of the zinc finger like motif across diferent species from human to sea sponge**. The alignment was carried out using Clustal Omega (EMBL-EBI).The conserved MBL family motifs are labeled as 1–4 and the cannocial β-CASP motifs are labeled A–C. The conserved residues that made up the novel zinc finger like structure are highlighted in green.

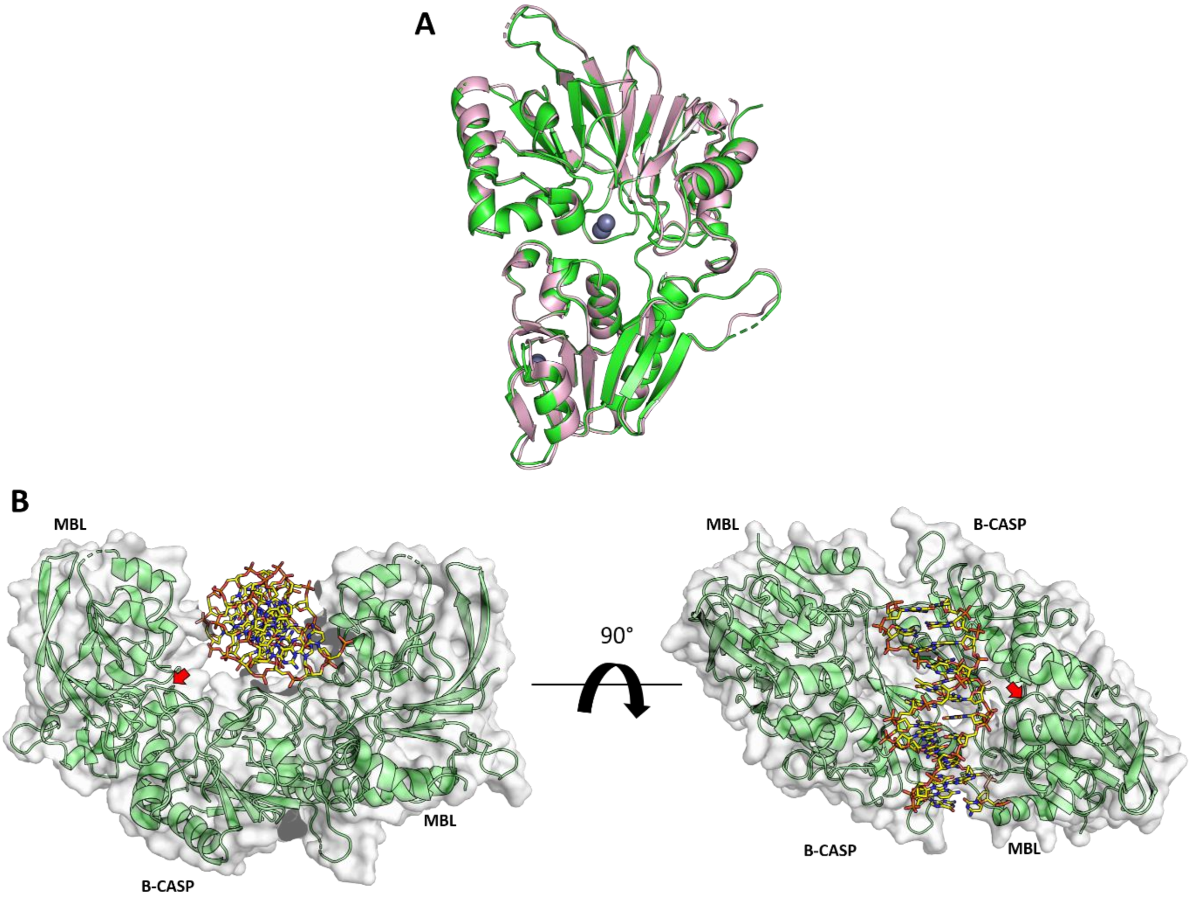


**Supplementary Figure 4: Structural features of Karim *et. al.,* 6WNL A:** Overlay of our Artemis structure 7AF1 in pink and 6WNL in green. **B:** Re-anaylsis of 6WNL with a model of DNA hairpin (yellow) refined in the solvent chanel. The active site of both Artemis molecules are indicated with the red arrows.

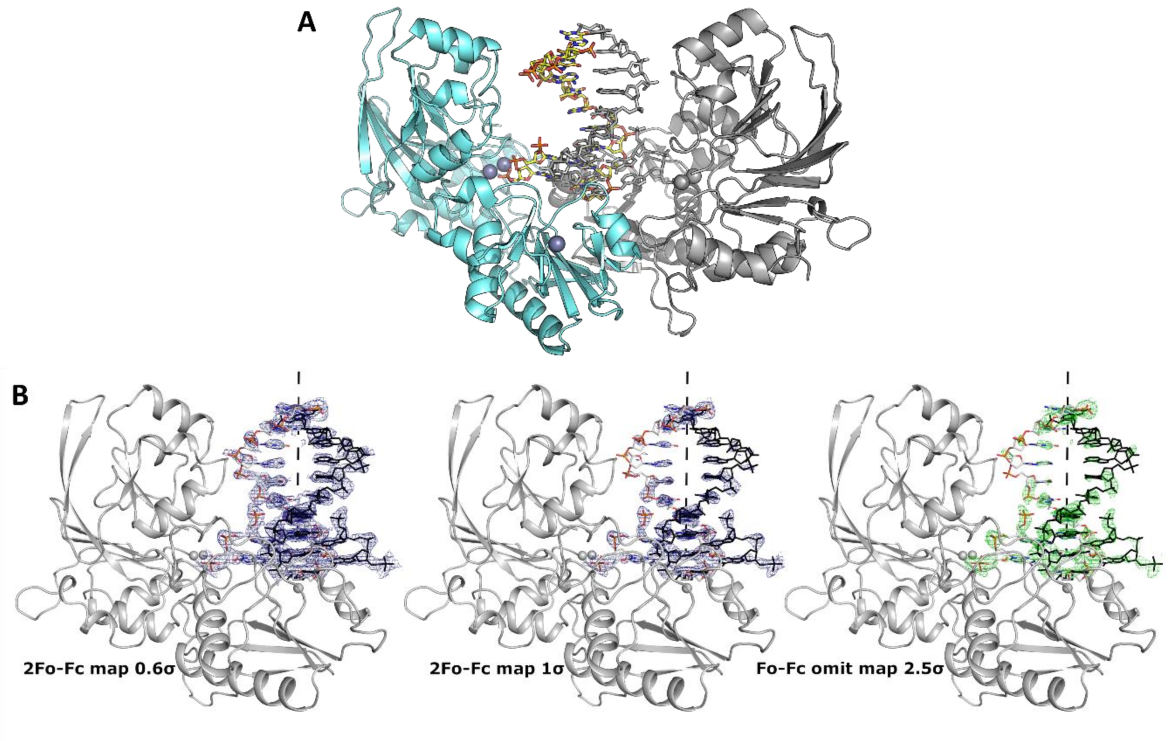


**Supplementary Figure 5: Overall strcuture of DNA bound Artemis, PDB code: 7ABS. A:** The structure of DNA bound Artemis showing a two-fold symmetry axis, where the symmetry molecule is shown in gray. **B:** The structure of DNA bound Artemis with diferent map contour.

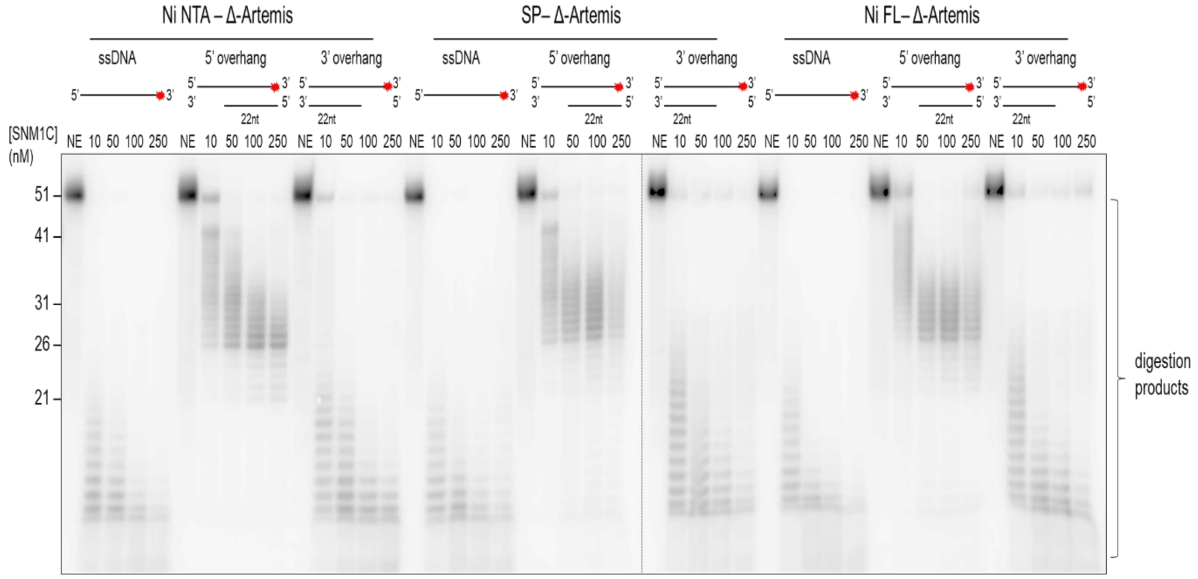


**Supplementary Figure 6: Nuclease assay of WT Artemis**. Full-length and truncated Artemis constructs purified in two diferent ways. Increasing amount of enzyme (concentrations as indicated), was incubated with 10 nM of either ssDNA, 5’ overhang, or 3’ overhang DNA substrate for 45 min at 37 °C.

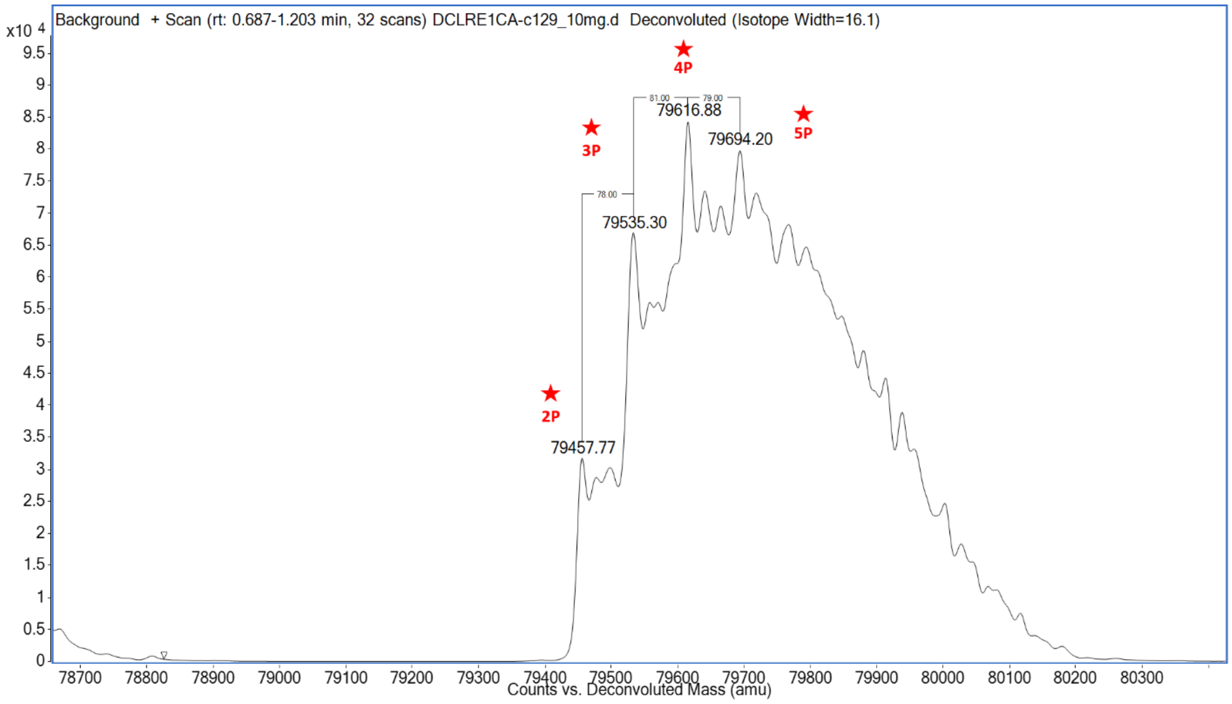


**Supplementary Figure 7: Intact mass analysis of the full length Artemis (aa 1–693) showing five different phosphorylation state (red stars) of the protein.**

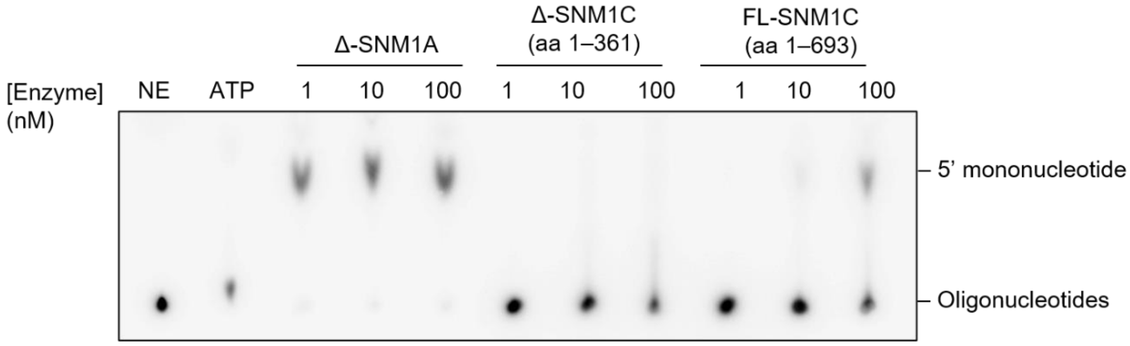


**Supplementary Figure 8: Full-length Artemis exhibits weak exonuclease activity, and Δ-Artemis negligible exonuclease activity.** Exonuclease activity examine by thin layer chromatography utilising increasing enzyme concentrations (as indicated) incubated with 10 nM of 5'-labelled ssDNA (50 nt) for 20 minutes at 37 °C. The samples were spotted on PEI-cellulose TLC sheets and developed with 400 mM phosphate buffer, pH 4.3 as the liquid phase. ATP and oligonucleotides remain near the origin, while the 5'-mononucleotide released by exonuclease activity migrates with the liquid phase.

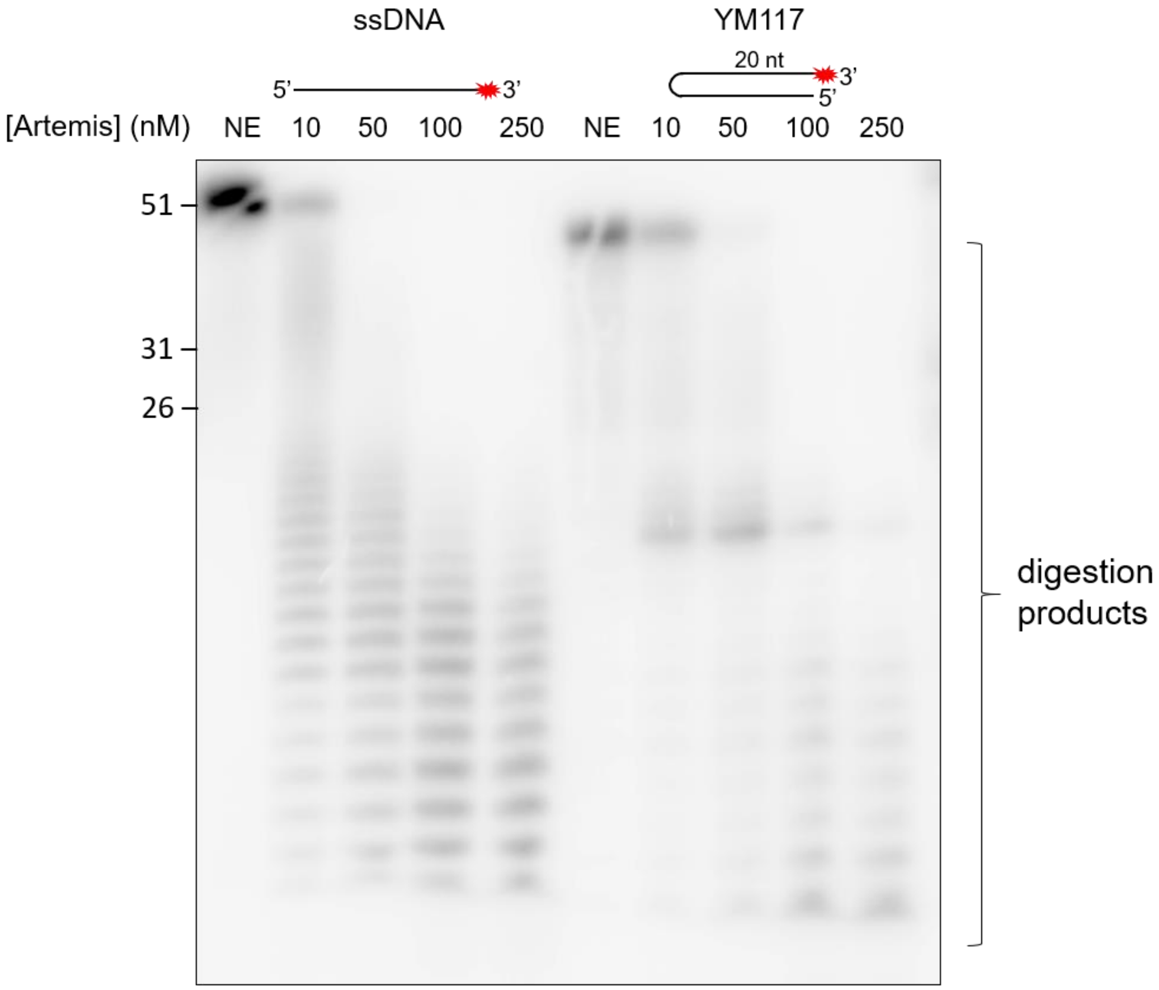


**Supplementary Figure 9: WT Artemis exhibits hairpin opening activity.** Increasing amounts (from 0 to 250 nM) of WT Artemis was incubated with 10 nM of 51 nucleotide ssDNA substrate or a duplex hairpin substrate (YM117 from Ma *et al.,* 2002) for 45 min at 37°C. Reaction products were subsequently analysed by 20% denaturing PAGE. The sizes (in nucleotides) of the marker oligonucleotides are indicated on the left-hand side of the corresponding bands.

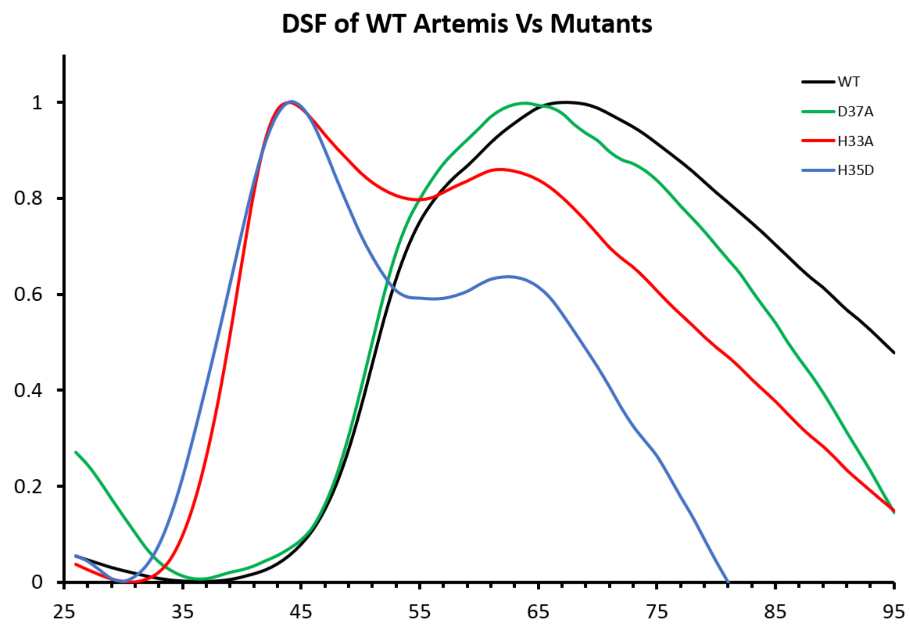


| | Tm |
|---|---|
| **WT** | 52 |
| **H35D** | 38 |
| **H33A** | 39 |
| **D37A** | 51 |

**Supplementary Figure 10: Thermal stability of Artemis WT and mutants assayed using differential scanning fluorimetry (DSF).** The H35D (blue) and H33A (red) variants exhibit a 13–14°C lower melting temperature than the wild-type protein, whilst the D37A mutation (green) barely affects the denaturation curve. The melting temperature of the proteins are presented in the right hand side table.

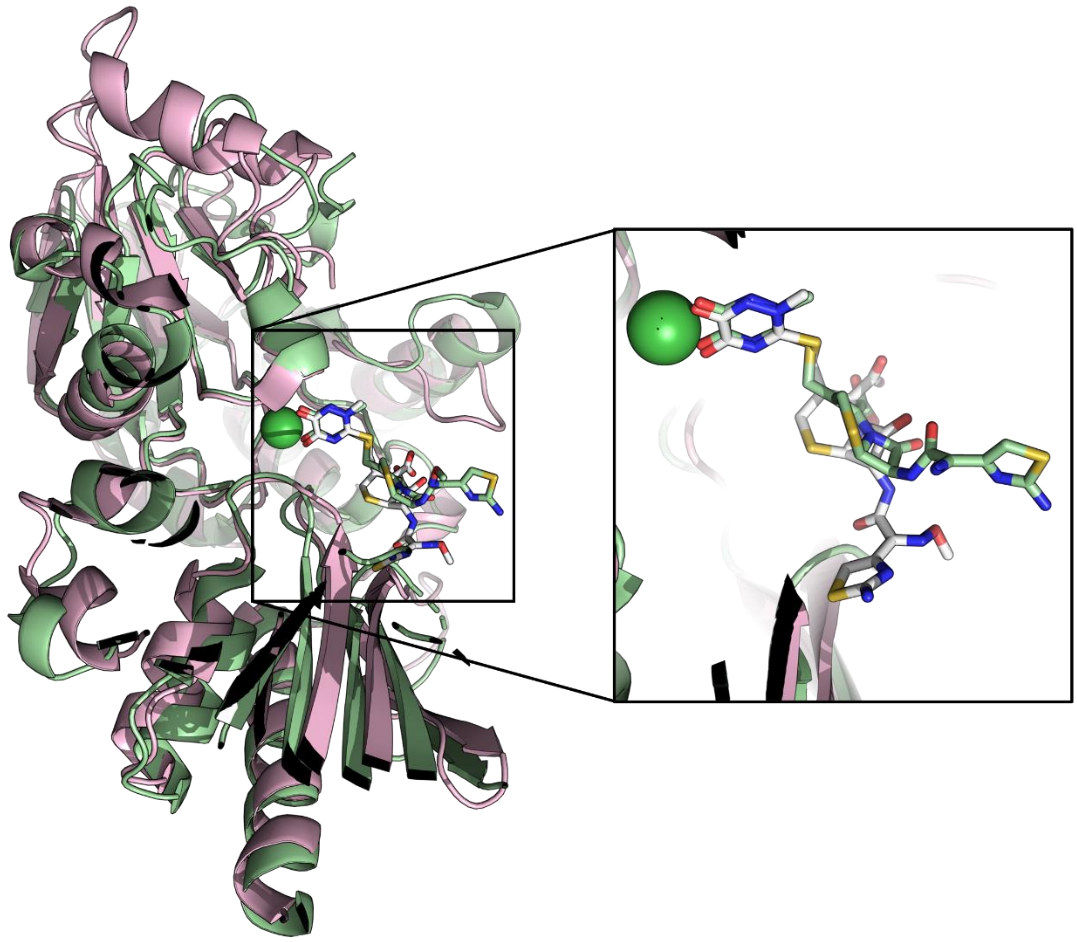

**Supplementary Figure 11: Structures of SNM1A and SNM1C (Artemis) in complex with ceftriaxone.** An overlay of SNM1A in green and SNM1C in pink, both in complex with ceftriaxone. An inset showing the binding mode of ceftriaxone to the metal centre through the cyclic 1, 2 diamide functional group of the compound.

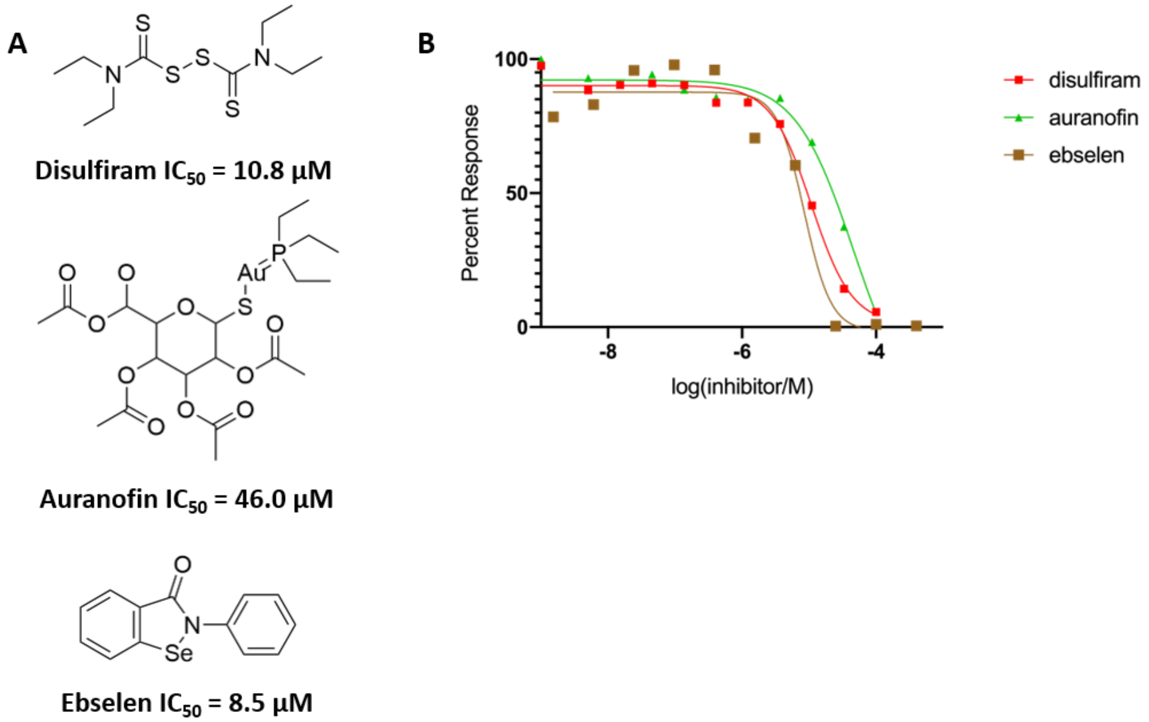


**Supplementary Figure 12: Inhibition of truncated Artemis (aa 1–361) by compounds containing thio-reactive groups** **A:** Molecular structure of disulfiram, auranofin and ebselen, and their coresponding $IC_{50}$ values. **B:** Inhibition curves of the three compounds against Artemis obtained using the real-time fluorescence-based nuclease assay.

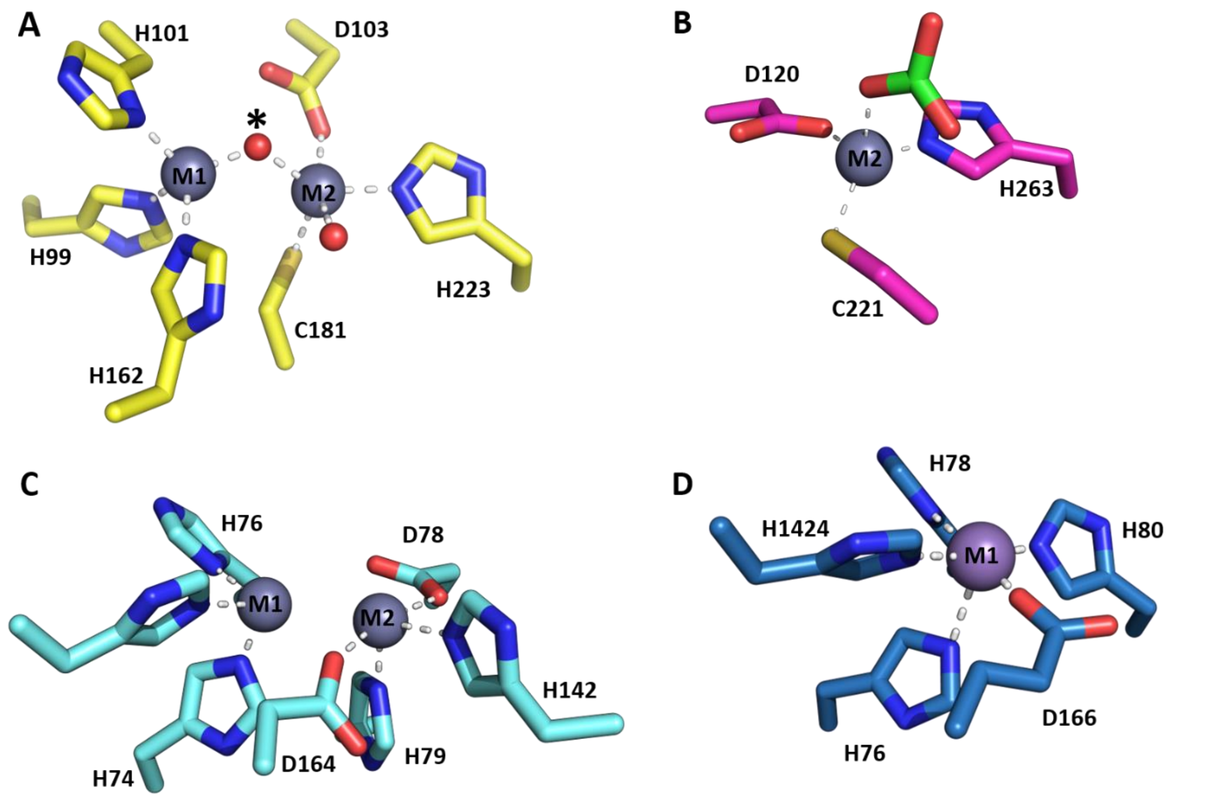


**Supplementary Figure 13: Active site views of bacterial MBL and bacterial MBL/ β-CASP enzymes A:** The active site of bacterial metallo-β-lactamase (subfamily B1) from *Bacteroides fragilis* (PDB: 1ZNB) coordinating 2 ZN (in grey). A water molecule shared (asterisk*) between the two metals is the proposed nucleophile for the hydrolytic reaction. **B:** The active site of Carbapenemase CphA from Aeromonas Hydrophylabacterial metallo-β-lactamase subfamily B2 (PDB: 1X8G) coordinating a mono zinc metal ion (in grey) with acetate (green) bound in the active site. **C:** The active site structure of RNase J1 from *Bacillus subtilis* (PDB: 3ZQ4 )containing di zinc (in grey). **D:** The mono metal active site structure of RNase J2 from *Staphylococcus epidermidis* (PDB: 6K6W) containing a single manganese ion (in purple).

| Code | DNA oligonucleotide sequence |
|---|---|
| **1A** | 5’ P-ATA AAT ATT TTT TAT TAA TAA TAG ATC ACC TTT CTT TCT CTT CTC CCC TT-OH 3’ |
| **1B** | 5’ OH-AT AAA TAT TTT TTA TTA ATA ATA GAT CAC CTT TCT TTC TCT TCT CCC CTT-OH 3’ |
| **1C** | 5’ biotin-AT AAA TAT TTT TTA TTA ATA ATA GAT CAC CTT TCT TTC TCT TCT CCC CTT-OH 3’ |
| **2** | 5’ OH-AAG GGG AGA AGA GAA AGA AAG GTG ATC TAT TAT TAA TAA AAA ATA TTT AT-OH 3’ |
| **3** | 5’ OH-AAG GGG AGA AGA GAA AGA AAG G-OH 3’ |
| **4** | 5’ OH-ATT ATT AAT AAA AAA TAT TTA T -OH 3’ |
| **5** | 5’ OH-TTC CCC TCC TCT CCT TCC TTC CTG ATC TAT TAT TAA TAA AAA ATA TTT AT-OH 3’ |
| **6** | 5’ OH-AAG GGG AGA AGA GAA AGA AAG G-OH 3’ |
| **7** | 5′ OH-GAT TAC TAC GGT AGT AGC TAC GTA GCT CTA CCG TAG TAA T-OH 3′ |
| **8** | 5’ OH-[FITC]TAA TTA ATA ATA GAT CAC CT [BHQ1]-OH 3’ |

| 3’ labelled substrates (α-$^{32}$P-dATP) | | | |
|---|---|---|---|
| **Annealed DNA Sequences** | **Substrate Structure** | **Description** | **Figure(s)** |
| 1A* | 5’ PHO 3’ | Single-stranded 51 nt DNA with a 5’ phosphate | 7, 9, Suppl. 6 |
| 1B* | 5’ OH 3’ | Single-stranded 51 nt DNA with a 5’ hydroxyl | 7 |
| 1C* | 5’ BIO 3’ | Single-stranded 51 nt DNA with a 5’ biotin | 7 |
| 1A* + 2 | 5’ 3’ / 3’ 5’ | dsDNA | 7 |
| 1A* + 3 | 5’ PHO 3’ 22 nt | 5’ overhang with the labelled strand bearing a 5’ phosphate | 7, Suppl. 6 |
| 1B* + 3 | 5’ OH 3’ 22 nt | 5’ overhang with the labelled strand bearing a 5’ hydroxyl | 7 |
| 1C* + 3 | 5’ BIO 3’ 22 nt | 5’ overhang with the labelled strand bearing a 5’ biotin | 7 |
| 1A* + 4 | 5’ PHO 3’ 22 nt | 3’ overhang with the labelled strand bearing a 5’ phosphate | 7, Suppl. 6 |
| 1B* + 4 | 5’ OH 3’ 22 nt | 3’ overhang with the labelled strand bearing a 5’ hydroxyl | 7 |
| 1C* + 4 | 5’ BIO 3’ 22 nt | 3’ overhang with the labelled strand bearing a 5’ biotin | 7 |
| 1A* + 5 | 22nt 3’ 5’ 3’ 5’ | Splayed arm | 7 |
| 1A* + 5 + 3 | 22nt 3’ 5’ 3’ 5’ | Leading strand flap | 7 |
| 1A* + 5 + 6 | 22nt 3’ 5’ 3’ 5’ | Lagging strand flap | 7 |
| 1A* + 3 + 5 + 6 | 22nt 3’ 5’ 3’ 5’ | Replication fork | 7 |
| 7* | 20 nt 3’ 5’ | 20 nt hairpin substrate (YM117 from Ma *et al.*, 2002) | Suppl. 9 |
| **5’ labelled substrates (γ-$^{32}$P-dATP)** | | | |
| 1B* | 5’ 3’ | Single stranded 50 nt DNA labelled with a γ-$^{32}$P-dATP | Suppl. 8 |
| **Fluorescently labelled substrates (used for inhibitor screening)** | | | |
| 8 | 5’ 3’ | 20 nt ssDNA substrate with a 5’ FITC and a 3’ BHQ-1 | 11, Suppl.12 |

**Supplementary Table 1: List of DNA oligonucleotide sequences used to generate simple and more complex DNA substrates.**

(a): the top panel indicates each oligonucleotide sequence as numbered.

(b): to generate substrates with the indicated structures, single-stranded oligos were annealed as described in the Materials and Methods. The codes used to generate each structure are as described. The red asterisk indicates the location of the radiolabel; the yellow asterisk, the position of a FITC moiety; and a dark blue circle, the location of a BHQ-1.
